# Activation of the Crtc2/Creb1 transcriptional network in skeletal muscle enhances weight loss during intermittent fasting

**DOI:** 10.1101/2020.06.27.175323

**Authors:** Nelson E. Bruno, Jerome C. Nwachukwu, Sathish Srinivasan, Richard Hawkins, David Sturgill, Gordon L. Hager, Stephen Hurst, Shey-Shing Sheu, Michael D. Conkright, Kendall W. Nettles

## Abstract

The Creb-Regulated Transcriptional Coactivator (Crtc) family of transcriptional coregulators drive Creb1-mediated transcription effects on metabolism in many tissues, but the in vivo effects of Crtc2/Creb1 transcription on skeletal muscle metabolism are not known. Skeletal muscle-specific overexpression of Crtc2 (Crtc 2 mice) induced greater mitochondrial activity, metabolic flux capacity for both carbohydrates and fats, improved glucose tolerance and insulin sensitivity, and increased oxidative capacity, supported by upregulation of key metabolic genes. Crtc2 overexpression led to greater weight loss during alternate day fasting (ADF), selective loss of fat rather than lean mass, supported by maintenance of higher energy expenditure during the fast. ADF downregulated most of the mitochondrial electron transport genes, and other regulators of mitochondrial function, that were substantially reversed by Crtc2-driven transcription. Glucocorticoids acted with AMPK to drive atrophy and mitophagy, which was reversed by Crtc2/Creb1 signaling. Crtc2/Creb1-mediated signaling coordinates metabolic adaptations in skeletal muscle that explain how Crtc2/Creb1 regulates metabolism and weight loss.

## 1. Introduction

Long-term dieting is usually not effective in maintaining a lower body weight due to increased hunger and a slower metabolic rate (1). In humans, loss of >10% of body weight also increases metabolic efficiency, such that ~500 less calories per day are required to maintain the same energy expenditure (2). Weight loss reduces the metabolic rate due to a number of changes including increased circulating leptin and decreased thyroid hormone and sympathetic nervous system (SNS) activity (2–4). People who maintain long-term weight loss almost invariably exercise (5), and both preclinical and clinical data suggest that this is through raising metabolic rate (6–8), decreased energy efficiency (9), and increased energy expenditure from the increased physical activity (10).

Creb1-regulated transcription is stimulated by GPCRs, such as the β-adrenergic receptor (βAR) during exercise (11–14), which activate adenylyl cyclases to convert ATP to the second messenger cAMP, which in turn liberates the catalytic subunit of protein kinase A (PKA) (15). Crtc1–3 family members are primary coactivators for Creb1 that are also highly conserved in the cAMP transcriptional response. Crtc1–3 are rendered cytoplasmic and inactive by phosphorylation (16). In contrast, they are fully activated through synergistic dephosphorylation by phosphatases downstream of PKA and calcium flux in polarizable tissues including the skeletal muscle (11), hippocampus (17), cortex (18), islet cells (16), and the renal proximal tubule (19). In skeletal muscle, Crtc/Creb1 is activated by exercise downstream of the β-adrenergic receptor and calcium flux (11), stimulating anabolic transcriptional programs (20), including induction of PGC-1a (11, 21) and in vitro enhancement of oxidative capacity (21). While Crtc/Creb1 coordinates metabolic transcriptional programs in other tissues such as adipocytes, liver, and pancreas, its role in skeletal muscle metabolism has not been studied in vivo.

To investigate the effects of Crtc2/Creb1 activation in adult skeletal muscle, we generated doxycycline (dox)-inducible transgenic mice that show skeletal muscle-specific overexpression of Crtc2 (FLAG-Crtc2 S171A/S275A/K628R mutant that prevents cytoplasmic localization) under the control of a tetracycline response element (11). Induction of Crtc2 improved whole animal and skeletal muscle metabolism, enabling maintenance of higher energy expenditure during fasting and improved weight loss.

## 2. Materials and Methods

### 2.1 Statistical analyses

Changes in body weight and body were analyzed with 2-way analysis of variance. For VO_2_, respiratory quotient, and ambulation data, the ad libitum and fasted periods were analyzed separately with 2-way ANOVA, and the refeeding data analyzed with Student’s T-test. For the energy consumption data, the fasted and re-fed data were analyzed together with 2-way ANOVA. Gene expression data were analyzed with 1-way ANOVA, with Dunnett’s test for multiple comparisons of each gene to its corresponding control. Glycemia and insulin changes over time were analyzed with 2-way ANOVA, and the AUC assessed with Student’s T-test. The behavioral data for meal consumption were analyzed with Student’s T-tests.

### 2.2 Crtc2 transgenic mice and animal care

All animal procedures were approved by the Scripps Research Institute ICACUC. TRE-Crtc2^tm^ mice were generated by cloning FLAG-Crtc2 (S171A, S275A, K628R) into pTRE-Tight. Oocyte injections were conducted into C57Bl6 at the University of Cincinnati. Double transgenic mice (DTg) were generated by crossing C57BL/6-Tg(TRE-Crtc2tm)1Mdc/j mice with 1256[3Emut] MCK-rtTA transgenic mice, which expresses the reverse tetracycline transactivator (rtTA) regulated by the 1256[3Emut] MCK promoter specifically in skeletal muscle (Grill *et al,* 2003). Wild type and TRE littermates were used as controls and treated with DOX in the same manner as DTg mice. High fat diet (Research Diet D12450J) contained 60 kcal% fat and matched control low fat diet (D12492) contained 10kCal% fat and 7% sucrose.

### 2.3 Glucose and insulin tolerance tests

Glucose tolerance testing was performed after an overnight fast. Blood glucose was quantified at 15, 30, 60, 90 and 120 min post-i.p. administration of 20% D-glucose (1 g/kg fasted body weight) with an Accu-check glucose monitor as described (Heikkinen *et al*, 2007). Insulin tolerance testing was performed on 6 hr-fasted mice injected i.p. with human insulin (1.5 U/kg fasted body weight). Blood glucose levels were determined immediately before, and 15 and 30 min after, injection.

### 2.4 In vivo electroporation

Plasmid DNA (50 mg) was transfected into mouse tibialis anterior (TA) muscles by electroporation as described previously (11). 1 hr before electroporation mice were anesthetized, and a small incision was made through the skin covering the TA muscle and then injected with 30 Al of 0.5 U/Al hyaluronidase and then injected with plasmid DNA. *Crtc2* and *GFP* expression plasmids were purified from DH5a *Escherichia coli* cultures using an EndoFree plasmid kit (Qiagen, Valencia, CA, USA), and resuspended in 71 mM sterile PBS. After the injections, electric pulses were administered by using non-invasive platinum-coated tweezertrodes. Eight 100-ms square-wave electric pulses at a frequency of 1 Hz and at 200-ms intervals were delivered with an ECM 830 electroporation unit (BTX-Harvard Apparatus, Holliston, MA, USA) with a field strength of 40 V/cm. After the electroporation procedure, the incision was closed with Vetbond surgical glue.

### 2.5 Indirect calorimetry and energy expenditure

Mice were individually housed in a sixteen chamber Oxymax comprehensive lab animal monitoring system (CLAMS; Columbus Instruments, Columbus OH) at 26° C. Animals were allowed to acclimate for 1 day followed by VO_2_ and VCO_2_ were measured at 13 min intervals over a continuous duration of 60 hr. Energy expenditure (EE) was calculated using the Lusk equation, kcal/h = (3.815 + 1.232 × RQ) × VO_2_ in l/h, where RQ is the ratio of VCO_2_ to VO_2_. Opto-Varimetrix-3 sensor system in the *x-, y-,* and *z*-axes of each chamber recorded sterotype, ambulatory and *z*-axis activity. Consecutive adjacent infrared beam breaks in either the *x*- or *y*-axes were scored as ambulatory activity.

### 2.6 Body composition

Body composition was determined using time domain nuclear magnetic resonance (TD-NMR) in a Bruker minispec.

### 2.7 Automated food intake monitoring

Feeding behavior was assessed using a BioDAQ episodic Food Intake Monitor (BioDAQ, Research Diets, Inc) as described (Stengel *et al,* 2010). Feeding parameters were calculated with BioDAQ Monitoring Software 2.2.02. Feeding bouts were defined as a change in stable weight of food. Meals consisted of two or more bouts within 5 s with a minimum meal amount of 0.02 g. Clusters of bouts occurring greater than 600 seconds apart were considered separate meals. Water or water containing Dox was provided *ad libitum* from regular water bottles. Mice habituated to the new environment within 1 day and showed normal food intake and regular body weight gain.

### 2.8 mRNA-seq and bioinformatics

Total RNA was extracted from tissues using TRIzol™ reagent (Invitrogen), and cleaned up using the RNeasy kit with on-column DNA digestion (QIAGEN). Three mRNA-seq libraries per condition were prepared using the NEBNext Ultra II directional RNA library Prep kit for Illumina (New England Biolabs), and read in paired-end 40-cycle sequencing runs on the NextSeq 500 system (Illumina). Sequences were aligned to the Ensembl *Mus musculus* GRCm38.83.chr.gtf genome (mm10) assembly using HISAT2 v2.0.5 (PMID: 25751142), sorted and indexed using SAMtools v1.7 (22) and quantified for differential expression analysis using Subread v1.6.1 featureCounts (23). Differential expression analysis was performed using DESeq2 v1.18.1 (24, 25). Enriched canonical pathways were identified using the Ingenuity pathways analysis software (QIAGEN). Gene Ontology (GO) analysis excluding inferred electronic annotations, was performed using the Mouse Genome Informatics GO Browser and GO Term Mapper (http://www.informatics.jax.org). Genes encoding mouse nuclear hormone receptors were obtained from the Nuclear Receptor Signaling Atlas (NURSA) (26). Genes encoding mitochondrial proteins were obtained from the mouse Mitocarta2.0 (27, 28). Genes encoding transcriptional coregulators were obtained from the EpiFactors database (29). Glucocorticoid-responsive genes were obtained by combining previously published datasets (30) reanalyzed with GEO2R, with other dexamethasone-regulated genes that we identified in C_2_C_12_ myotubes, GEO accession no. GSE149453. Heatmaps were prepared using Pheatmap v1.0.8, RColorBrewer v1.1–2, and other software packages available in R. Other data visualizations were performed using GraphPad Prism7.

### 2.9 Glucose uptake

Insulin mediated 2-deoxy-D-[^14^C] glucose uptake was determined by incubating primary myotubes with PBS or PBS containing 100 nM insulin for 30 min followed by addition of 0.1 mM of cold 2-deoxy-D-glucose and 0.1μCi/well of 2-deoxy-[^14^C] D-glucose for 5 min prior to solubilization. Nonspecific deoxyglucose uptake was measured in the presence of 20 μM cytochalasin B and subtracted from the total glucose uptake.

### 2.10 Primary myoblast isolation and culture

Primary myoblasts were isolated from P1 to P3 (1–3 day old) C57BL/6J pups as described (Springer *et al,* 1997) and cultured on collagen coated plates in media consisting in 1:1 F-10/DMEM supplemented with 20% FCS and 25 μg/ml bFGF. Subconfluent myoblasts were differentiated into multinucleated myotubes using a 1:1 F-10/5 mM glucose DMEM supplemented with 4% heat inactivated horse serum (Hyclone) for up to five days. For steroid-free culture conditions, cells were grown in charcoal:dextran-stripped FBS (Gemini Bioproducts, cat no. 100-119).

### 2.11 Adenoviruses

Control GFP-expressing and Crtc2 expressing adenoviruses have been described (Koo *et al*, 2005; Wu *et al,* 2006). For all experiments myotubes were infected after 72 hr of differentiation with 4×10^8^ viral particles per ml per well for 24–48 hr.

### 2.12 Immunoblotting analysis

Immunoblot analysis was performed as described (Conkright *et al,* 2003a). Whole cell extracts were prepared in HEPES whole cell lysis buffer (20 mM HEPES, pH 7.4, 1% TX-100,1x phosphoSTOP complete inhibition cocktail tablets [Roche], and 1X Complete-EDTA Free protease inhibitor cocktail [Roche]). Frozen muscle tissue was homogenized in lysis buffer T-PER^®^ Tissue Protein Extraction reagent (Thermo Scientific) by a polytron homogenizer and sonication. The protein concentration was determined using the protein assay reagent (Bio-Rad). 60–80 μg of protein were separated by SDS-PAGE and transferred to nitrocellulose membranes for immunoblotting. Nur77/Nr4a1 #2518 Rabbit polyclonal antibody was from Novus Biological. CRTC2 #A300-638A Rabbit polyclonal antibody was from Bethyl Laboratories. Irs2 (L1326) #3089S Rabbit polyclonal antibody was Cell Signaling Technology, Inc. Tubulin #T9026-.2ML Mouse Monoclonal antibody was from Sigma-Aldrich. pAKT(Thr308) (244f1)Rb and pAKT(Ser473) (971s) were from Cell Signaling Technology. PGC1α ab54481 Rb was from Abcam.

### 2.13 Quantitative RT-PCR (qPCR)

Total RNA from tissue or cells was harvested using Rneasy RNA Kit (Qiagen) and cDNA prepared using Transcriptor High Fidelity cDNA Synthesis Kit (Roche). Relative abundance of cDNAs was determined a Roche LightCycler 480, and the resulting data were normalized to *Rpl-23* ribosomal RNA (IDT) as described (Conkright *et al,* 2003b). The primer pairs are as follows: *Nr4a1* (5’-ttctgctcaggcctggtact’, 5’-gattctgcagctcttccacc-3’); *Nr4a3* (5’-tcagcctttttggagctgtt’, 5’-tgaagtcgatgcaggacaag-3’); *Irs2* (5’-acaacctatcgtggcacctc-3’, 5’-gacggtggtggtagaggaaa-3’); Idh3a (5’-gtgacaagaggttttgctggt-3’, 5’-tgaaatttctgggccaattc-3’); *Cytc* (5’-gatgccaacaagaacaaaggt-3’, 5’-tgggattttccaaatactccat-3’); *Glut4* (5’-tgtggctgtgccatcttg-3’, 5’-cagggccaatctcaaagaag-3’).

### 2.14 Mitochondrial DNA (mt-DNA) analysis

Total DNA was extracted using the GenElute™ mammalian genomic DNA miniprep kit (Sigma-Aldrich). Triplicate real-time PCR reactions were performed using 50 ng of total DNA, *CoxI* (5’-agcattcccacgaataaataaca-3’, 5’-agcattcccacgaataaataacat-3’) or *CoxII* (5’-tttcaacttggcttacaagacg-3’, 5’-tttcaacttggcttacaagacg-3’) mitochondrial DNA primers, and a control genomic DNA primer, *Gapdh* (5’-gaaaaggagattgctacg-3’, 5’-gcaagaggctaggggc-3’).

### 2.15 Histology

Dissected muscle tissue was mounted in OCT medium (TissueTek) and froze in liquid N2– cooled isopentane. Tissue sections were stained for succinate dehydrogenase activity to distinguish between oxidative and nonoxidative fibers using standard methods.

### 2.16 Fatty acid metabolism

Fatty acid oxidation was assayed by incubating (9,10(n)-^3^H) palmitic acid (60 Ci/mmol) bound to fatty-acid free albumin (final concentration:100 μM, palmitate:albumin 2:1) and 1 mM carnitine with primary myocytes for 2 hr. Tritiated water released was collected and quantitated.

### 2.17 High-content imaging and analysis

Primary or C_2_C_12_ myotubes in black 96- or 384-well tissue culture plates with clear bases (Greiner Bio-One, North America, Inc.) were stained for 15 min with 200 nM MitoTracker^®^ Orange CM-H2TMRos dye (Invitrogen™ by ThermoFisher Scientific). The myotubes were rinsed with steroid-free DMEM to remove un-incorporated dye, and then incubated in the differentiation media for 45 min to reach peak fluorescence. Myotubes were fixed in 4% formaldehyde for 20 min, stained with 300 nM DAPI for 5 min, permeabilized in PBS containing 0.1% Triton X-100 for 20 min, blocked for 1 h with 1x TBS containing 0.1% Tween-20 (TBS-T) and 2.5% normal goat serum, and incubated at 4°C overnight with an AlexaFluor^®^ 488-conjugated anti-skeletal muscle myosin (F59) antibody (Santa Cruz Biotechnology, Inc. cat no. sc-32732 AF488). The next day, the myotubes were washed 4 times with TBS-T to remove unbound antibodies and rinsed twice with PBS. The stained myotubes were imaged at 10–20X magnification on the IN Cell Analyzer 6000 platform (GE Healthcare). For each treatment condition, a stack of 24–27 images containing an average of 50–100 myotubes per image were analyzed using the IN Cell Developer Toolbox image analysis software, with a customized segmentation protocol for myotubes. The average (mean) diameter and mitochondrial potential (i.e. MitoTracker staining density x area) of myotubes in these images were then calculated.

## 3. Results

### 3.1 Crtc2 stimulates mitochondrial activity and improves oxidative capacity

After one week of dox treatment, Crtc2 mice showed increased mitochondrial succinate dehydrogenase activity in the gastrocnemius muscle (**Figure 1A, Supplemental Fig. 1A–B**) while myotubes overexpressing Crtc2 or Crtc3 oxidized more palmitate than control myotubes (**Figure 1B**), as previously reported (21). The Crtc2 mice displayed increased expression of *Cytochrome C (Cycs), Ppargc1a/PGC-1a* (as previously reported (11, 21), and *Nr4a3/NOR-1* nuclear receptor (**Figure 1C**) which induces mitochondrial biogenesis and increases oxidative capacity (31, 32), and is induced by high intensity exercise (11). Crtc2 also upregulated the *Dgat1* gene (**Figure 1C**), which is a direct Creb1 target gene (11), the product of which catalyzes a rate-limiting step that increases IMTG stores. These mitochondria-related genes were also upregulated in primary myotubes overexpressing Crtc2, indicating a skeletal muscle-intrinsic effect of Crtc2 that mimics the effects of exercise (**Figure 1D**). We also observed a doubling of mitochondrial DNA after in vivo electroporation of Crtc2 into skeletal muscle (**Figure 1E**). These data support a model where exercise induced mitochondrial biogenesis uses Crtc2/PGC-1a transcriptional control programs to upregulate mitochondrial biogenesis and oxidative capacity.

**Figure 1.**
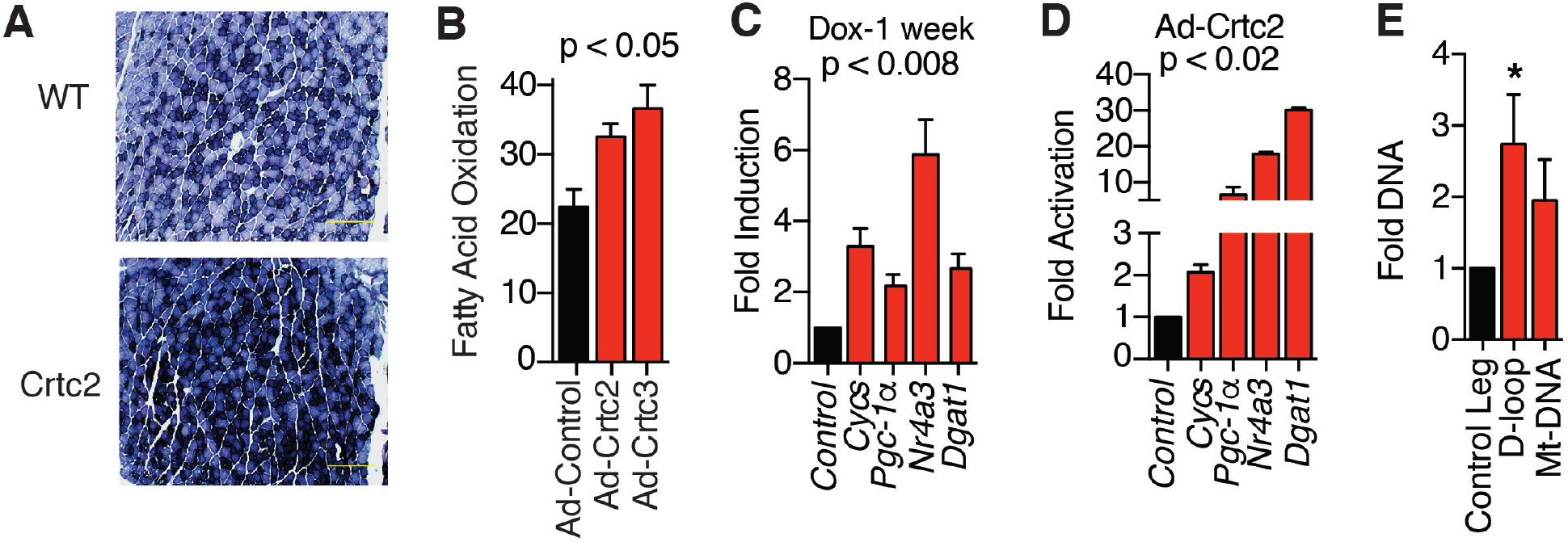
Crtc2 expression in skeletal muscle enhances mitochondrial biogenesis. (**A**) Histological analysis of succinate dehydrogenase in myofibers from gastrocnemius muscle sections after dox treatment. Scale bar = 500 μm. (**B**) Oxidation of ^3^H-Palmitic acid in the supernatant of primary myocytes transduced with control, Crtc2, or Crtc3-expressing adenovirus. (**C**) Gene induction calculated relative to WT mice. *N* = 8 mice per group. (**D**) Gene expression in primary myocytes transduced with adenovirus expressing Crtc2 or GFP control 72 hr post-transduction. (**E**) Crtc2 expression plasmid was electroporated into the tibialis anterior muscle and GFP plasmid in the contralateral leg. After 7 days, DNA was extracted for assessment of mitochondrial DNA (mt-DNA) for Crtc2/GFP control. *N* = 5 mice per group. * p < 0.05. Data are shown as mean + SEM.

### 3.2 Crtc2 improves glucose tolerance and carbohydrate metabolism

To probe for Crtc2-dependent changes in carbohydrate metabolism, we performed glucose tolerance tests comparing Crtc2 and WT mice. After one week of dox treatment, the Crtc2 mice displayed significantly improved glucose disposal, including a dramatic improvement in their ability to reduce glycemia (**Figure 2A, Supplemental Figure 2**), and required a lower insulin excursion (**Figure 2B**). Crtc2 overexpression also improved insulin tolerance (**Figure 2C**). In primary myotubes, expression of Crtc2 was sufficient to significantly increase the uptake of 2-Deoxy-D-glucose (**Figure 2D**).

**Figure 2.**
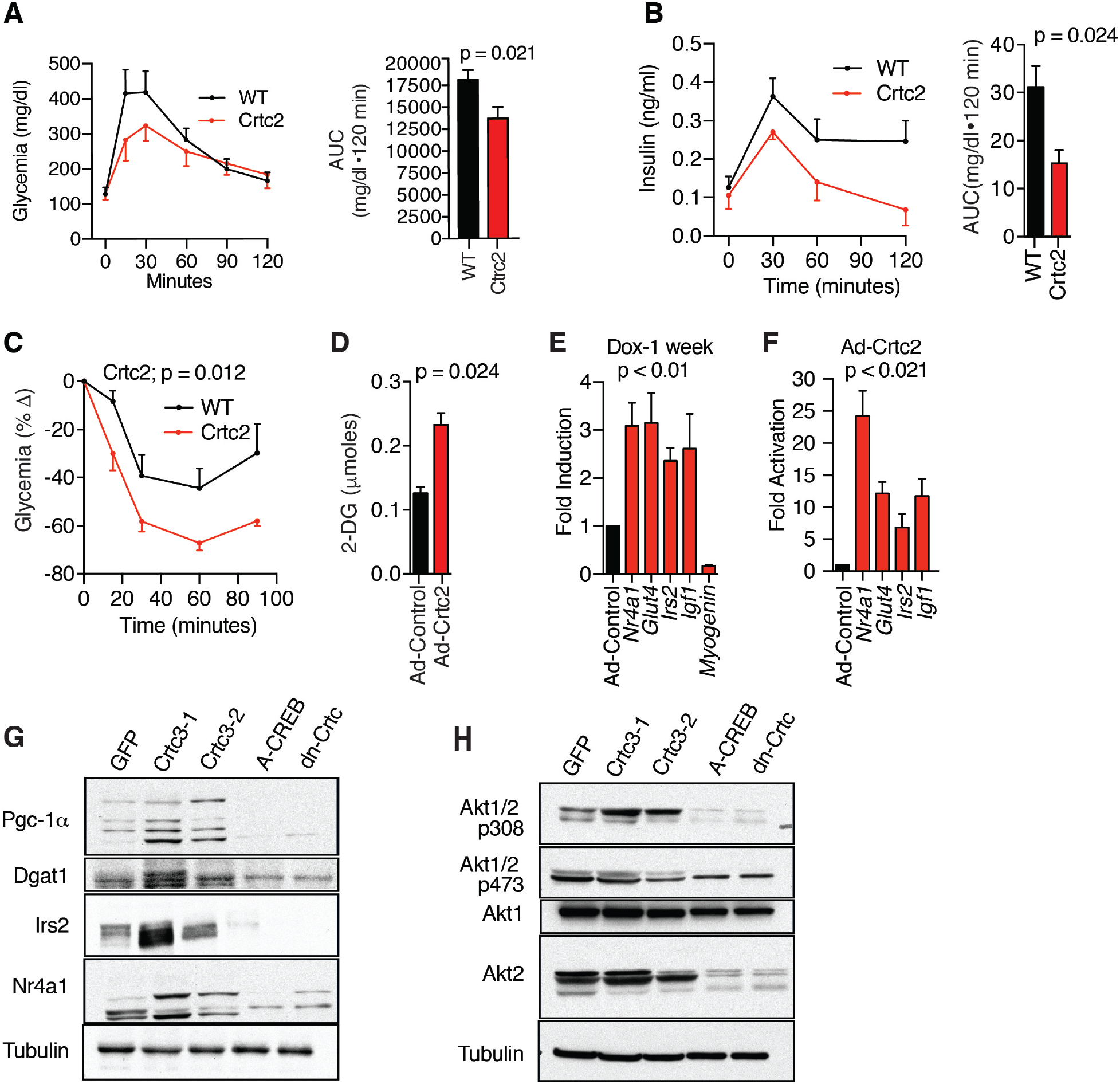
Crtc2 transgenic mice maintain enhanced glucose disposal and insulin tolerance. (**A**) Glucose tolerance test (GTT). Right panel, area under the curve. *N* = 10 mice per group (**B**) Plasma insulin levels during GTT described in panel **B**. (**C**) Insulin tolerance tests. *N* = 10 mice per group (**D**) Insulin-mediated uptake of 2-deoxy-D-glucose in primary myocytes transduced with adenovirus expressing Crtc2 or GFP control. *N* = 10 per group. Student’s t-test. (**E**) Crtc2 mice were treated **+/-** dox for 1 week and the gene induction compared by qPCR relative to the no dox groups. N= 10 per group. (**F**) Gene expression in primary myocytes transduced with adenovirus expressing Crtc2 or GFP control were compared by qPCR. *N*=4 per group. (**G–H**) Myocytes stably expressing GFP, Crtc3 (lanes 2 and 3), dominant negative Creb (A-CREB), or dominant negative Crtc2 (dn-Crtc) were compared by western blots. Data are mean ± SEM.

Crtc2 mice and Crtc2/3-overexpressing myotubes upregulated a number of carbohydrate signaling genes in normal fed conditions at both the RNA and protein levels (**Figure 2E–H**), including the glucose transporter, *Glut4,* whose insulin-dependent translocation to the membrane drives glucose uptake into skeletal muscle, accounting for 80% of glucose disposal. The *Nr4a1* gene encodes Nur77, a nuclear receptor family transcription factor that regulates transcriptional programs for carbohydrate metabolism, lipolysis, and mitochondrial biogenesis (33), and is upregulated by high intensity exercise or adrenergic signaling (11). *Irs2, Akt1,* and *Akt2* encode core components of the insulin receptor signaling pathway that can coordinate anabolic signaling and improve carbohydrate metabolism. We also identified a significant inhibition of myogenin expression in the normal fed Crtc2 mice (**Figure 2E**), a primary driver of muscle atrophy during fasting. To validate on target mechanism of action, we generated stable cell lines overexpressing Crtc3, dominant negative A-Creb, or dominant negative Crtc2 expressing only the Creb binding domain (16). These results demonstrate that A-Creb and dn-Crtc2 blocked basal expression of PGC-1a, Irs2, and Akt2, and prevented phosphorylation of Akt1/2 at Thr308 (**Figure 2G–H**). These experiments demonstrate that exercise coordinates beneficial metabolic responses that are recapitulated by activation of skeletal muscle Crtc2/Creb1 transcriptional programs.

### 3.3 Crtc2 overexpression in skeletal muscle facilitates weight loss

We noticed during the GTTs that the Crtc2 mice tended to lose more weight after the overnight fast, so we tested them in the context of a common dieting scheme of intermittent fasting. We studied aged Crtc2 (rtTA/Crtc2^tm^) or WT (rtTA/WT) mice, which in the C57BL/6 background naturally gain fat mass as they age (18 weeks, 30–39 g, **Figure 3A**). During dox treatment, Crtc2 mice gained weight similarly to control WT mice (**Supplemental Fig. 3A**). When subject to alternate day fasting (ADF), three separate cohorts of Crtc2 mice displayed significantly greater and sustained weight loss than WT mice, with Crtc2 mice losing about twice as much weight in each case, 9-14% of body weight (**Figure 3B, Supplemental Fig. 3B**).

**Figure 3.**
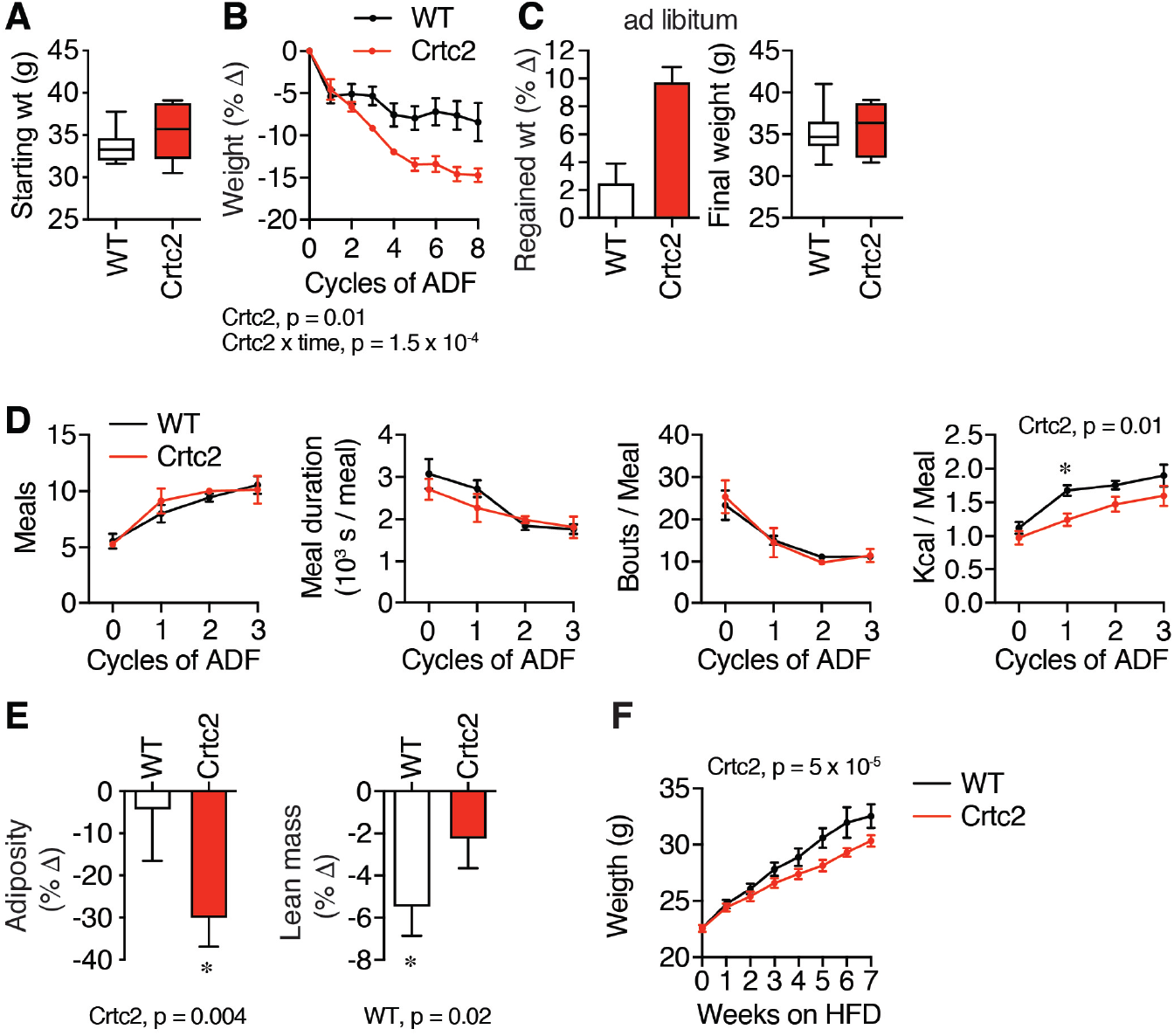
Crtc2/Creb1 signaling enables greater loss of weight and adiposity during alternate day fasting (ADF). (**A**) 18-week old C57BL/6 male control (WT) and Crtc2 mice were treated with dox for 35 days and weighed. Data are shown as box-and-whisker plot, indicating the middle quartiles and range. *N* = 6 (WT) or 5 (Crtc2) mice per group. (**B-E**) Mice from (**A**) were subject to alternate day fasting (ADF). (**B**) Changes in the body weight during ADF. (**C**) Weight regain after the ADF treatment and 3 weeks of ad libitum feeding. (**D**) Behaviors were assayed with Biodak monitoring system, including number of meals, bouts per meal, meal duration, and kilocalories consumed over the 12-hr re-feeding period. *significant in Sidak’s test for planned comparisons. (**E**) Adiposity and lean mass determined by whole-body NMR measurements taken before and after 8 cycles of ADF. Data analyzed with 1-way ANOVA (**F**) 8-week old C57BL/6 male control and Crtc2 mice were treated with dox and placed on a high fat diet (HFD). *N* = 9 (WT) or 7 (Crtc2) mice per group Time course data was analyzed by 2-way ANOVA. Data are shown as mean ± SEM.

After 3 weeks of ad libitum refeeding, the Crtc2 mice re-gained more weight to catch up with the WT mice (**Figure 3C**), which may be due to the anabolic effects of higher insulin sensitivity (**Figure 2**). We found that locomotion was similarly reduced in both groups during the refeeding period (**Supplemental Fig. 3C**), the number of meals doubled in both groups, while meal duration and number of bouts per meal declined similarly (**Figure 3D**). We observed a reduction in kilocalories consumed in the Crtc2 relative to the WT group, which planned comparisons demonstrated was significant only during the second ADF cycle, when the WT mice increased consumption at a faster rate than the Crtc2 mice (**Figure 3D**). Increases in skeletal muscle storage of glycogen and triglycerides associated with the anabolic phenotype of Crtc2/Creb1 activation (11) likely contributed to the early response to fasting, perhaps explaining the initially slower rate of increased food consumption in the Crtc2 mice. These differences in caloric consumption are consistent with the differences in body weight between cohorts and disappear after normalization for body weight. However, control mice lost more lean mass, while Crtc2 mice conserved lean mass and burned the excess fat accumulated during aging (**Figure 3E**), similar to the effects of exercise (34). To verify that Crtc2 helps maintain lower body weight in a different model, we placed mice on a high fat diet, and found that Crtc2 mice also maintained lower body weight than WT mice in this context (**Figure 3F**). To test for pre-fasting changes in the central control of feeding and energy expenditure, we completed a leptin infusion study, as leptin secretion from adipocytes both suppresses hunger and stimulates the SNS. The WT and Crtc2 mice displayed identical patterns of weight loss following leptin infusion (**Supplemental Figure 4D**), and there were also no differences in plasma levels of feeding hormones, cholesterol, or triglycerides (**Supplemental Fig. 4E–F**). Thus, the activation of Crtc2/Creb1 signaling in skeletal muscle significantly modulated body composition and weight loss, mimicking the effects of exercise training including preferential loss of fat mass.

### 3.4 Crtc2 maintains energy expenditure during fasting

Lowered metabolic rate is one of the major defenses against fasting-induced weight loss. As expected, ad libitum oxygen consumption cycled with activity in the mice, which increased at night and decreased during the day (**Figure 4A, Supplemental Fig. 4A**). Fasting reduced oxygen consumption at every cycle, and this effect was more pronounced during the day as the fast was extended (**Figure 4A**). However, Crtc2 mice consistently showed higher oxygen consumption, independent of fasting (**Figure 4 A**).

**Figure 4.**
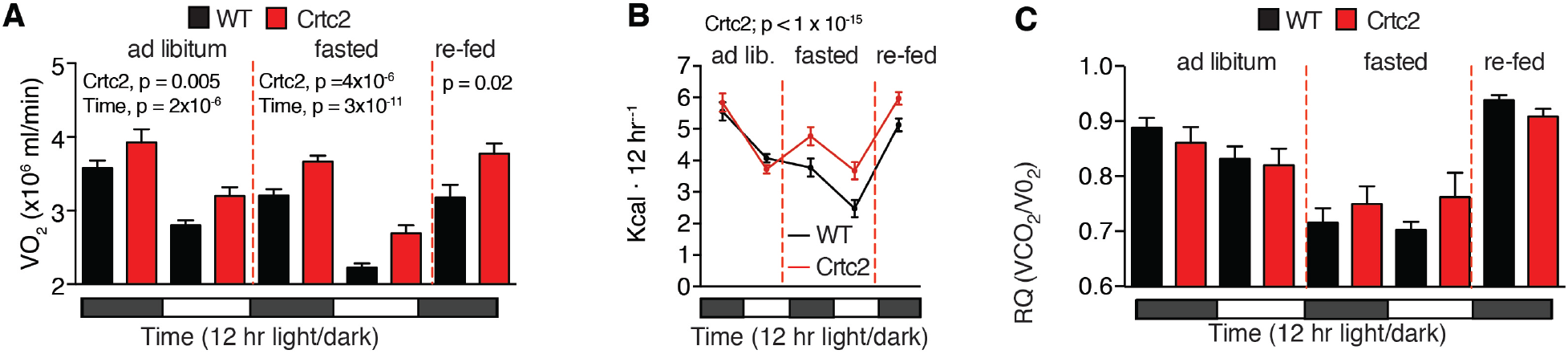
Whole-body energy expenditure is reduced by weight loss and fasting and is reversed by Crtc2 expression. (**A**) VO_2_ and VCO_2_ were measured continuously (**Figure S2A**) for 72 hrs. in a CLAMS animal monitoring system. Shown are the 12-hr averages. *N* = 8 mice per group. (**B**) Total energy expenditure was calculated from VO_2_ and VCO_2_ values using the Lusk equation. (**C**) The respiratory quotient (RQ) was determined during the CLAMS experiment. Data are shown as mean ± SEM and were analyzed by 2-way ANOVA for the different feeding conditions separately.

When fed ad libitum, Crtc2 and control mice display similar rates of energy expenditure (EE) (**Figure 4B**). We calculated EE from indirect calorimetry measurements of VO_2_ and VCO_2_ and performed ANCOVA analysis using body weight and NMR data as covariates to adjust the EE to total body weight and body composition (**Supplemental Fig. 4B**). While fasting, the EE of WT mice dropped significantly. In contrast, during fasting EE in Crtc2 mice did not drop below the ad libitum levels (**Figure 4B**). Respiratory quotient was also reduced during fasting, indicating greater utilization of fat for fuel, but the differences between WT and Crtc2 mice were not significant (**Figure 4C, Supplemental Fig. 4C**). Maintenance of a higher energy expenditure during fasting thus links Crtc2 activity to increased weight loss during fasting.

### 3.5 The transcriptional response to fasting and weight loss and its regulation by Crtc2

To understand the skeletal muscle intrinsic effects of Crtc2 on dieting and weight loss, we performed transcriptional profiling of mice tibialis anterior (TA) muscles electroporated with control and Crtc2 expression plasmids in contralateral legs. One cohort of electroporated mice was subject to 3 cycles of ADF, while the other cohort was fed ad libitum. TA muscle tissues were collected from both cohorts at the end of the third 24-hr fast. The transduction increased *Crtc2* mRNA levels by ~2.5-fold across groups (**Figure 5A**). In response to ADF, we identified 1,880 differentially expressed genes (DEGs) in the control-transduced muscle and 3,197 DEGs in Crtc2-transduced muscle, with 1,380 genes in common between the two groups (**Figure 5B**). Analysis of canonical signaling pathways and gene ontology (GO) annotations revealed that ADF modulated the transcriptional programs for lipid and carbohydrate metabolism, fatty acid beta-oxidation, and insulin signaling (**Figure 5C–F,** annotated gene lists are in **Supplemental Dataset1**). ADF also regulated transcriptional programs that control protein balance, including autophagy and ubiquitin-dependent proteasomal degradation, RNA splicing, and transcription (**Figure 5C-D**). There were also over 200 DEGs regulated by ADF that were annotated as transcription factors and transcriptional coregulators along with > 50 transcription factor and coregulators altered by Crtc2 over-expression (**Figure 3G–I**), demonstrating the complexity of the transcriptional response to intermittent fasting in skeletal muscle.

**Figure 5.**
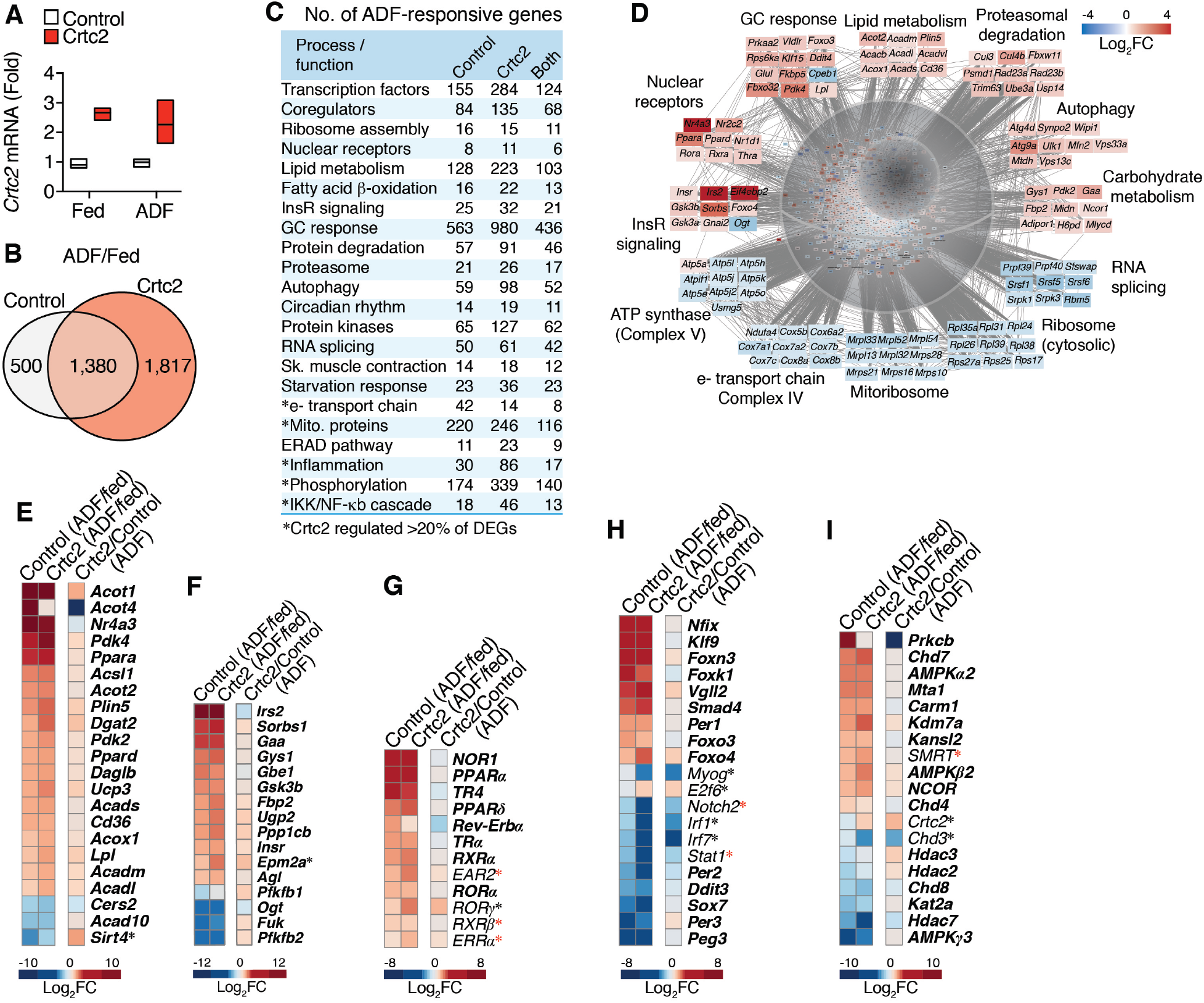
Transcriptional control of fasting and weight loss and its regulation by Crtc2. (**A**) Mice TA muscles were transduced with GFP control or Crtc2 expression vector. Crtc2 mRNA levels in the TA of mice fed ad libitum (Fed), or subject to ADF (Fast) were compared by qPCR. *N* = 3 mice per group. (**B**) Venn diagram showing the numbers of DEGs identified by mRNA-seq comparing the effects of ADF in control versus Crtc2-transduced TA muscles. (**C**) Biological processes and functions regulated by ADF and Crtc2. Gene ontology (GO) analysis suggests that ADF regulates several processes/ functions in a Crtc2-sensitive manner. The numbers of DEGs involved in each process are shown. See SI Appendix, **Dataset S1** for a complete list of represented GO annotations. (**D**) Examples of ADF-regulated transcriptional programs and mRNAs in control TA muscles. (**E–I**) Gene expression profiles in control and Crtc2-transduced TA muscles of mice subjected to ADF relative to the ad libitum fed mice (columns 1-2), and effect of Crtc2 transduction relative to control during ADF (column 3). Expression profiles of genes that encode **E**) nuclear receptors, **F**) other transcription factors, **G**) transcriptional coregulators, **H**) regulators of lipid metabolism, and **H**) regulators of carbohydrate metabolism. . ADF-regulated genes in control muscle appear in **bold**. *Crtc2-regulated genes in mice subjected to ADF. 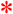ADF-regulated genes in Crtc2-transduced muscle but not control. FC, fold change.

We considered that Crtc2 might be regulating secreted proteins that modulate the response to dieting, using the Metazoan Secretome Knowledgebase (35). Among 2,500 genes encoding likely secreted proteins, 171 DEGs were regulated by Crtc2 during ADF (Secretome, **Supplemental Dataset1)**. Using the Reactome.org peer-reviewed pathway database, we detected significant enrichment of pathways regulating extracellular matrix, inter-cell signaling, and immune function, among others (**Supplemental Table 1**), but no changes in myokines that would drive weight loss. However, Crtc2 signaling in skeletal muscle would be expected to have effects on other tissues just from changes in skeletal muscle metabolic flux capacity (**Figures 1–2**).

### 3.6 Crtc2 reverses fasting-induced effects on mitochondria

In addition to lower metabolic rate, fasting and weight loss render humans and other mammals more energetically efficient, such that the same amount of work is performed with less ATP production, suggesting a remodeling of glycolytic or oxidative capacity (2, 36). ADF modulated the expression of >200 genes whose proteins localize in mitochondria according to MitoCarta (27) (**Figure 6A**). ADF downregulated most of the genes that encode the electron transport chain (complexes I–IV), the ATP synthase (complex V), the mitoribosome, and the cytosolic ribosome (**Figure 6A-C**, **Supplemental Fig. 5**), but significantly upregulated genes such as *Sdha, Uqcrc1,* and *Atp5a1* (**Figure 6B**, **Supplemental Fig. 5**).

**Figure 6.**
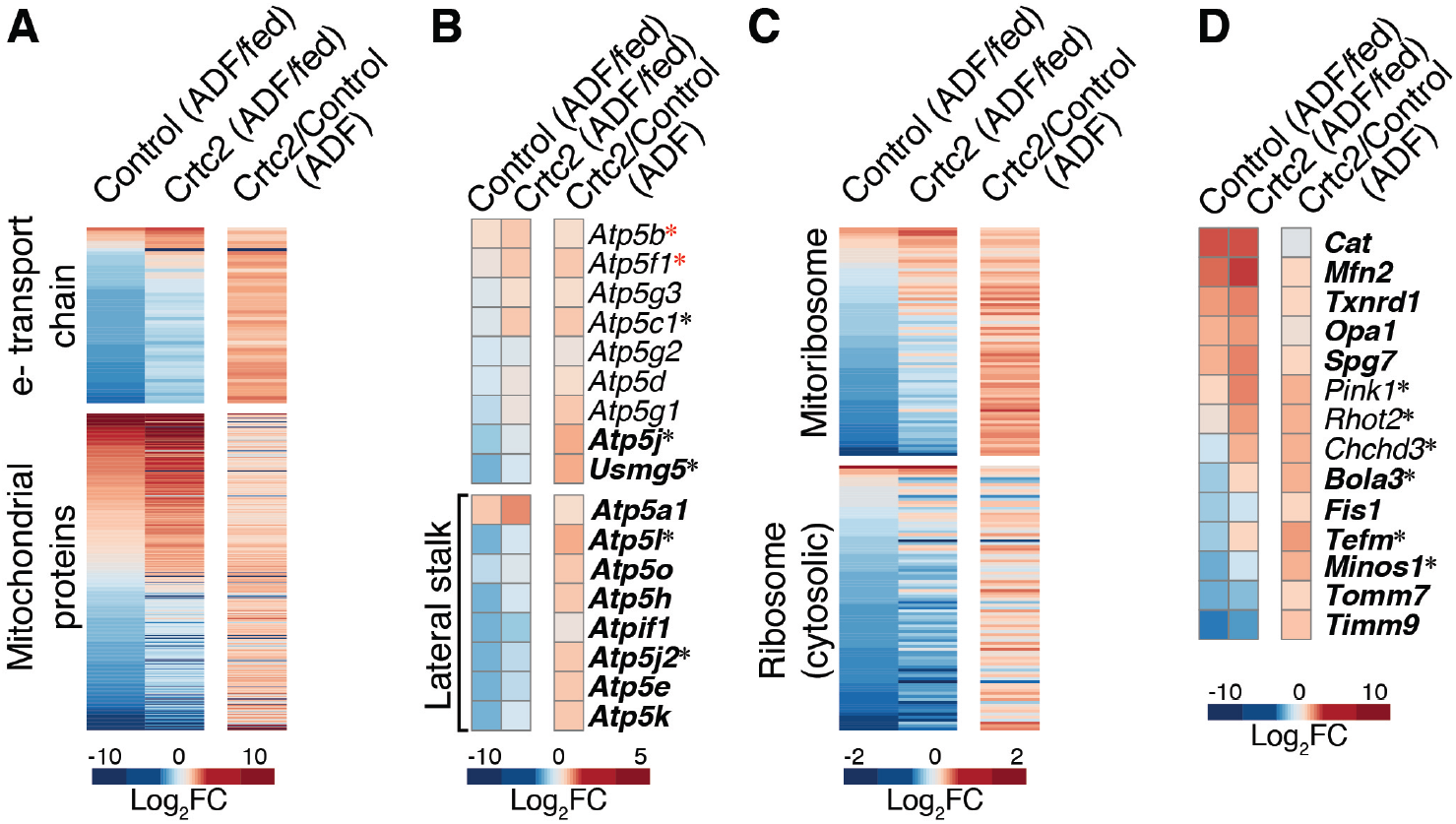
Alternate day fasting and Crtc2 regulation of mitochondrial gene expression. (**A–D**) Gene expression profiles in control and Crtc2-transduced TA muscles of mice subjected to ADF relative to the ad libitum fed mice (columns 1-2), and effect of Crtc2 transduction relative to control during ADF (column 3). The profiles of all genes encoding **A**) mitochondrial proteins, including subunits of electron transport chain (ETC) complexes, **B**) subunits of the mitochondrial ATPase (complex V), **C**) ribosomal subunits, and **D**) highlighted mitochondrial proteins, are shown. ADF-regulated genes in control muscle appear in **bold**. *Crtc2-regulated genes in mice subjected to ADF. 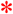ADF-regulated genes in Crtc2-transduced muscle but not control. FC, fold change.

We observed a significant ADF-induced expression of genes encoding ROS scavenger proteins such as thioredoxin *(Txnrd1)* and catalase (*Cat*), and genes regulating mitochondrial stability, *Tomm7, Timm9*, and the AAA protease *Spg,7* known to be active during the removal of damaged mitochondrial proteins (**Figure 6D**). Among the ATP synthase genes, there was a more pronounced loss of gene expression in the lateral stalk genes (**Figure 6B**), which may decrease the conductance of mitochondrial permeability transition pore (mPTP) and thus protect mitochondria from detrimental swelling and oxidative injury (37). ADF also induced increases in mitochondrial fusion *(Mfn2* and *Opa1),* and decreased fission *(Fis)* gene expression, consistent with the protective effects of fusion during nutrient deprivation (38). These adaptations could be advantageous for weight loss as they prepare the mitochondria for the reduced metabolic rate and increased reliance on lipid metabolism during fasting, without causing permanent damage to the mitochondrial pool within the cell.

In mice subjected to ADF, Crtc2 transduction altered expression of 80 genes whose proteins localize in the mitochondria. Crtc2 expression strongly attenuated the ADF-dependent repression of electron transport chain genes and the mitoribosome (**Figure 6A–C**, **Supplemental Fig. 5**), and generally upregulated the other genes encoding proteins that localize to mitochondria. In addition, Crtc2 enhanced expression of ribosomal genes that facilitate mitoribosome assembly, and genes such as *Spg7, Pink1, RhoT2,* and *Chchd3* that control the quality of mitochondrial proteins, but blocked ADF-dependent downregulation of *Minos1,* a central component of the mitochondrial inner membrane organizing system (**Figure 6D**). Crtc2 also upregulated expression of *Tefm,* the mitochondrial transcriptional elongation factor, and *Bola3,* which participates in electron transport chain assembly (39). These data suggest potential mechanisms to explain how physical activity—through Crtc2/Creb1-regulated transcriptional programs—can help maintain a higher metabolic rate and a lower body weight by upregulating expression of mitochondrial respiratory chain subunits and modulating expression of other mitochondria-localized gene products, consistent with the well-known beneficial effects of exercise on mitochondrial biogenesis and energy expenditure.

### 3.7 Crtc2 reverses catabolic effects of glucocorticoids on skeletal muscle

To further understand the signaling pathways that are regulated by fasting and Crtc2, we noted that glucocorticoids (GCs) are one of the primary drivers of systemic catabolism during fasting. In response to ADF and Crtc2 overexpression, we observed over a thousand DEGs that were annotated as “glucocorticoid response” genes (**Figure 5C**). In the context of an overnight fast, treatment with dexamethasone (Dex) more than doubled loss of lean mass (**Figure 7A**). Using high-content imaging, we observed that Dex reduced the diameter and mitochondrial potential of nutrient-deprived primary mouse skeletal myotubes (**Figure 7B**). GCs can also activate the master energy sensor, AMP-activated protein kinase (AMPK) (40), which can act as a glucocorticoid receptor (GR) coactivator (41). AMPK is activated by a lower ATP/ADP ratio during fasting to inhibit anabolism and stimulate catabolic processes such as autophagy and mitophagy. In primary myotubes, we observed a Dex-dependent activating phosphorylation of AMPK that increased over time, consistent with an underlying transcriptional mechanism (**Figure 7C**). Activation of AMPK with both Dex and the AMP analog, AICAR, had an additive detrimental effect on myotube diameter (**Figure 7D**), demonstrating that GR collaborates with AMPK to drive skeletal muscle atrophy. To show that glucocorticoids and Crtc2 are directly opposing each other in skeletal muscle, we transduced myotubes with Crtc2 expressing adenovirus, which increased tube diameter, effects that were reversed by treatment of the cells with Dex (**Figure 7E**). Importantly, the dominant negative A-Creb significantly reduced tube diameter, demonstrating that Crtc2 is acting through Creb1 to regulated anabolic effects.

**Figure 7.**
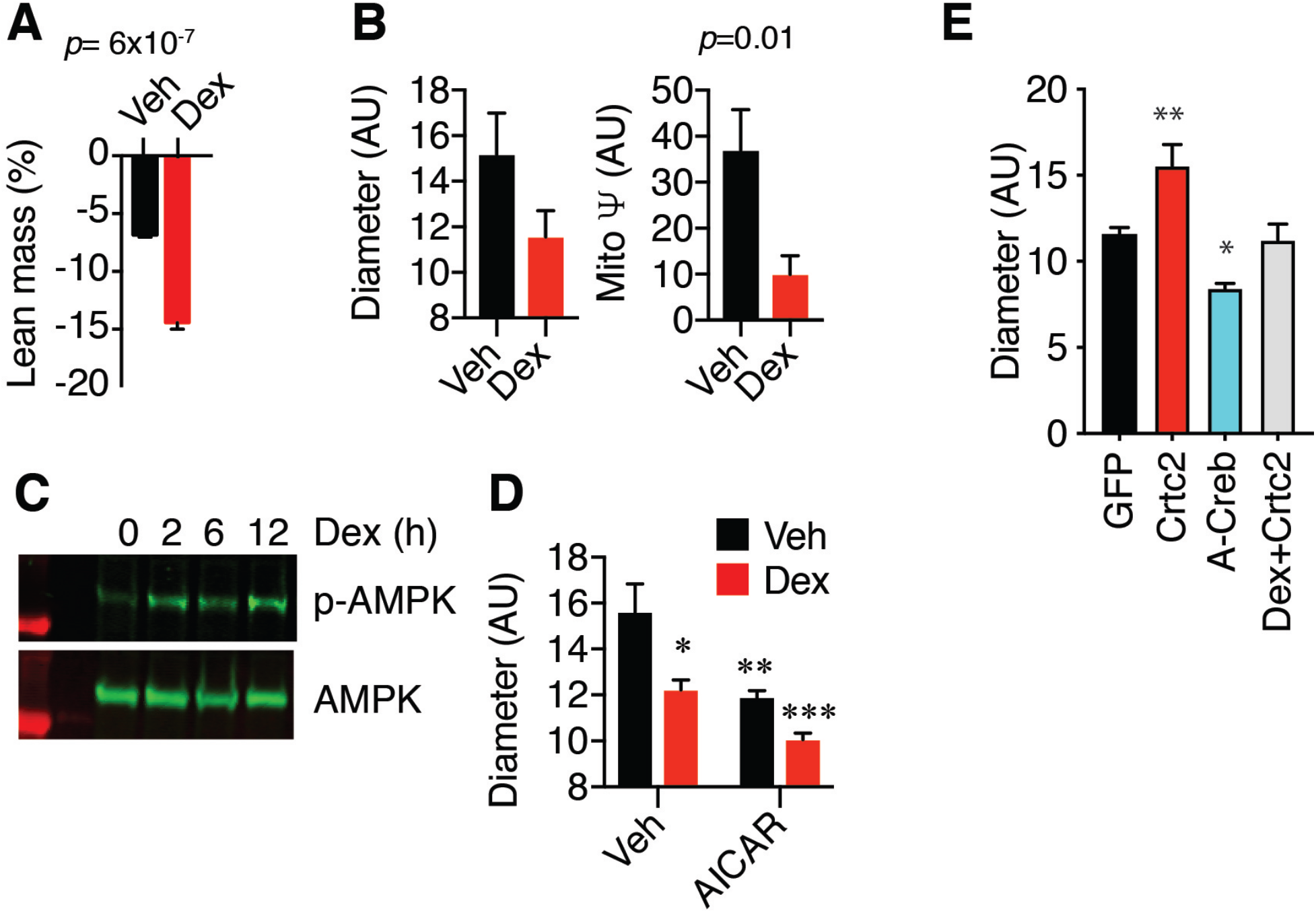
Glucocorticoids and AMPK regulate skeletal muscle atrophy and mitophagy. (**A**) Effects of Dex on lean body mass. Mice injected with vehicle or 10 mg/kg Dex were fasted overnight. Lean mass was measured by whole-body NMR before and after treatment. *N* = 5 mice per group. (**B**) Diameter and mitochondrial potential of steroid-deprived primary myotubes treated for 2 days with vehicle or 100 nM Dex were compared by confocal high-content imaging. *N* = 24-27 images x 50-100 myotubes per image. (**C**) Steroid-deprived C_2_C_12_ myotubes were stimulated with 100 nM Dex for 0-12 h. Levels of phosphorylated and total AMPK in whole lysates were compared by western blot. (**D**) Diameter of steroid deprived C_2_C_12_ myotubes treated for 48 h with vehicle, 100 nM Dex, or 1 uM AICAR were compared by confocal high content imaging. *N* = 24-27 images × 50-100 myotubes per image. (**E**) C_2_C_12_ myocytes were transduced with adenovirus GFP, Crtc2, or A-Creb genes and analyzed by confocal high content imaging. *N* = 24-27 images x 50-100 myotubes per image.

## Discussion

This work reveals mechanisms through which Crtc2-induced transcriptional programs dramatically rewire metabolism to regulate energy expenditure and the response to dieting. Crtc2 stimulated mitochondrial biogenesis and activity, upregulated β-oxidation genes, increased lipid flux capacity, improved insulin sensitivity and glucose disposal, and maintained higher energy expenditure during fasting, phenocopying metabolic effects of exercise. Overexpression of PGC-1a increases mitochondrial density and exercise performance, but without the increase in metabolic rate that can occur from exercise training (42–45), highlighting the requirement for additional exercised-induced signals. Transcriptional profiling revealed a profound effect of Crtc2 that prevented the ADF-dependent loss of mitochondrial electron transport chain gene expression and opposed catabolic effects of glucocorticoids that are induced by fasting. Crtc2/Creb1 metabolic rewiring enabled greater weight loss during dieting, providing a mechanism for how exercise facilitates weight loss.

This work adds to a body of evidence on Crtc2 and Creb1 as global regulators of metabolism in liver, pancreas, adipose tissue, and other endocrine tissues by coordinating with other tissue- and stressor-specific transcriptional signaling programs. We previously demonstrated that Creb1 and GR *work together* to coordinate gluconeogenesis transcriptional programs in the liver, where GR doubles the number of Creb1 binding sites and increases binding of Creb1 (46). We recently found that effects of GR ligands in skeletal muscle coordinately regulate inhibition of AKT and new protein synthesis, while stimulating mitophagy and protein degradation, but used different transcriptional pathways to inhibit insulin-mediated glucose uptake (accepted at *Nat Chem Biol,* posted on bioRxiv https://doi.org/10.1101/2020.06.15.153270). This work suggests that the effects of exercise on fasting may also involve a coordination of Creb1 and GR signaling as master transcriptional regulators of metabolism, in conjunction with other stress-response signals.

Reduced energy expenditure and increased hunger during fasting are partly mediated by a lowering of the satiety hormone leptin (2–4). Leptin is also secreted basally from adipose tissue in proportion to fat mass, so it is chronically lower during fasting, which lowers sympathetic tone and energy expenditure, thyroid hormone levels, and stimulating the HPA axis (47)(**Figure 8A**). Within skeletal muscle, increased glucocorticoids and AMPK activity are sufficient to reduce mitochondrial potential (**Figure 7)**, which along with likely contributions from fasting-induced lowering of thyroid hormone, bAR, and insulin receptor activity, induce mitochondrial dynamics and mitophagy (**Figure 8A**).

**Figure 8.**
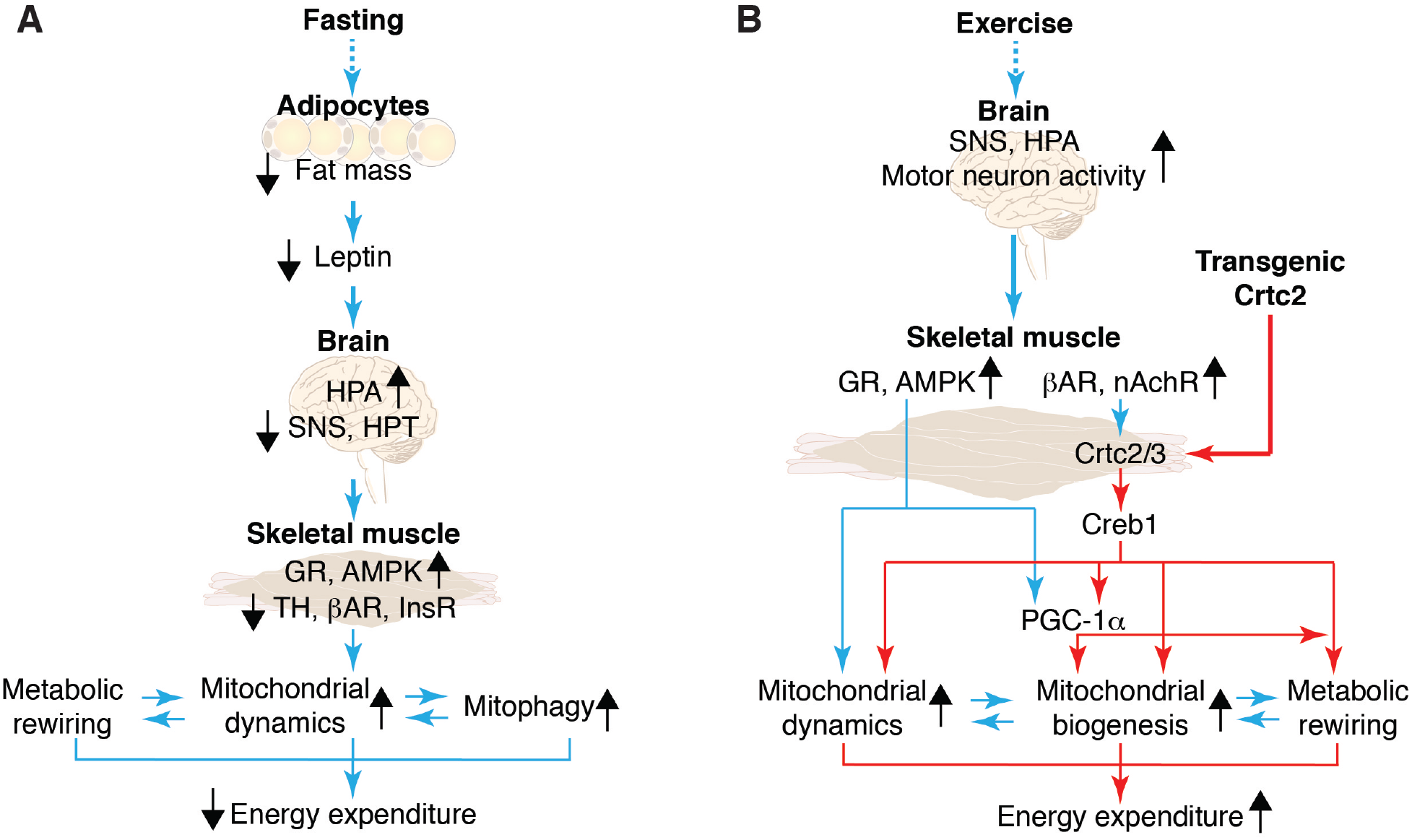
Hormonal and neural modulation of energy expenditure in skeletal muscle. (**A**) Fasting-induced regulation of mitochondrial dynamics and mitophagy. (**B**) Exercise-induced regulation of mitochondrial dynamics and mitochondrial biogenesis, which can be constitutively activated by overexpression of Crtc2 in skeletal muscle. bAR, b-adrenergic receptor; GR, glucocorticoid receptor; HPA, hypothalamic-pituitary-adrenal axis; HPT, hypothalamic-pituitary-thyroid axis; InsR, insulin receptor; nAchR, nicotinic acetylcholine receptor; SNS, sympathetic nervous system; TH, thyroid hormone

These energetic effects of fasting are reversed by exercise (**Figure 8B**), which we now show can be recapitulated by enforced expression of Crtc2/Creb1 and associated upregulation of mitochondrial biogenesis master regulatory genes, *Nr4a1, Nr4a3, Essrra/*ERRa, and *Ppargc1a/PGC-1a* (**Figure 1**). The combination of bAR and Ca^2+^ signaling are required to maximally stimulate nuclear translocation of Crtc2 in skeletal muscle (11). In this model, increased GC and AMPK activity during exercise promotes mitochondrial dynamics, which is then coupled with mitochondrial biogenesis to increase mitochondrial potential and mass, in coordination with other metabolic changes (**Figures 4–5**). Overexpression of Crtc2 circumvented the neuronal activation of Creb1, as Crtc2 mice had most of the hallmarks of exercise adaptations before fasting and weight loss (**Figures 4–5,** SI Appendix, **Figure S5B–C**), including increased exercise performance, myotube diameter, and storage of IMTG and glycogen (11). As the SNS is activated logarithmically in response to exercise intensity, Crtc2/Creb1 are likely driving responses to novel exercise, leading to both anabolic and metabolic reprogramming.

## Data Availability

The raw and processed RNA-seq dataset available at the National Center for Biotechnology Information (NCBI) Gene Expression Omnibus (GEO) https://www.ncbi.nlm.nih.gov/geo/ and can be retrieved with the accession number GSE149150, which will be released upon publication of this manuscript. The processed RNA-seq dataset is also provided in SI Appendix, **Dataset S1**. Relevant protocols and raw data not detailed in the Methods or Supplementary Information are available upon request.

## Acknowledgements

N.E.B. was supported by the BallenIsles Men’s Golf Association. J.C.N. was supported by the Frenchman’s Creek Women for Cancer Research.

## Author contributions

Conceptualization, N.E.B., M.D.C., K.W.N.; Methodology, N.E.B., M.D.C.; Investigation, N.E.B., J.C.N., S.S., R.H., D.S.; Writing - Original Draft, K.W.N., N.E.B., M.D.C., S.H., S.S.S.; Writing - Revision, K.W.N, J.C.N., N.E.B.; Supervision, M.D.C, G.L.H., K.W.N., S.S.S., N.E.B.

## Supplemental Information for

**Supplemental Table 1.**
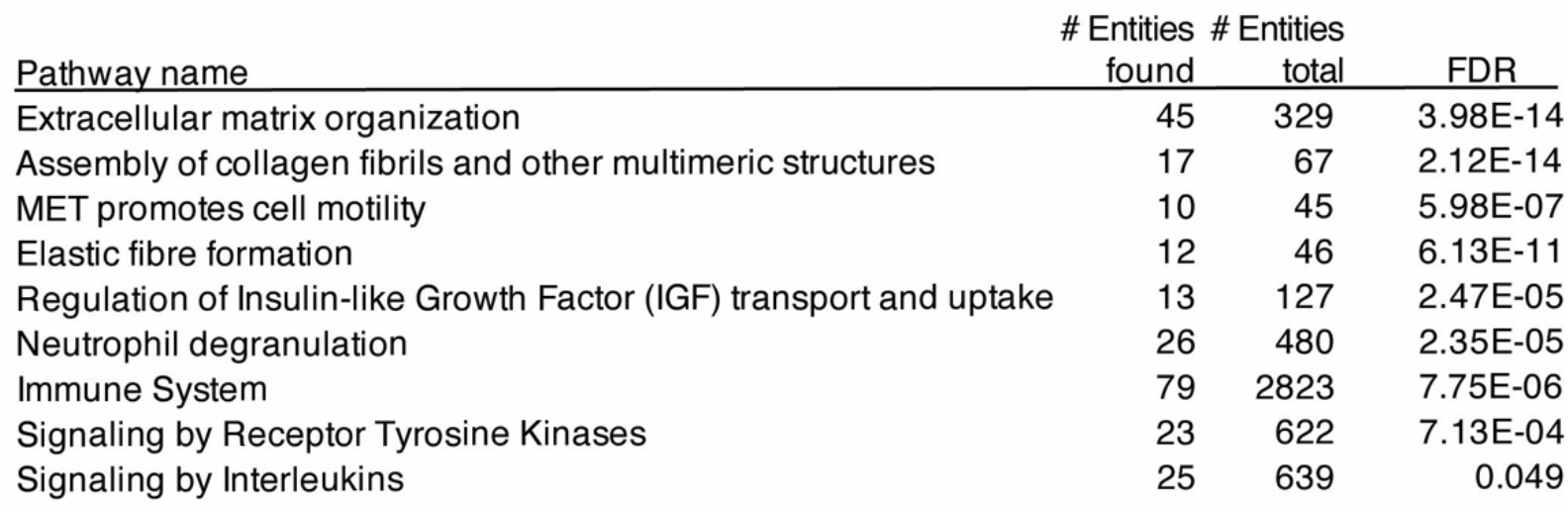
Gene pathway enrichment for secreted proteins among DEGs regulated by Crtc2 during ADF.

### Supplemental Figures

**Supplemental Fig 1.**
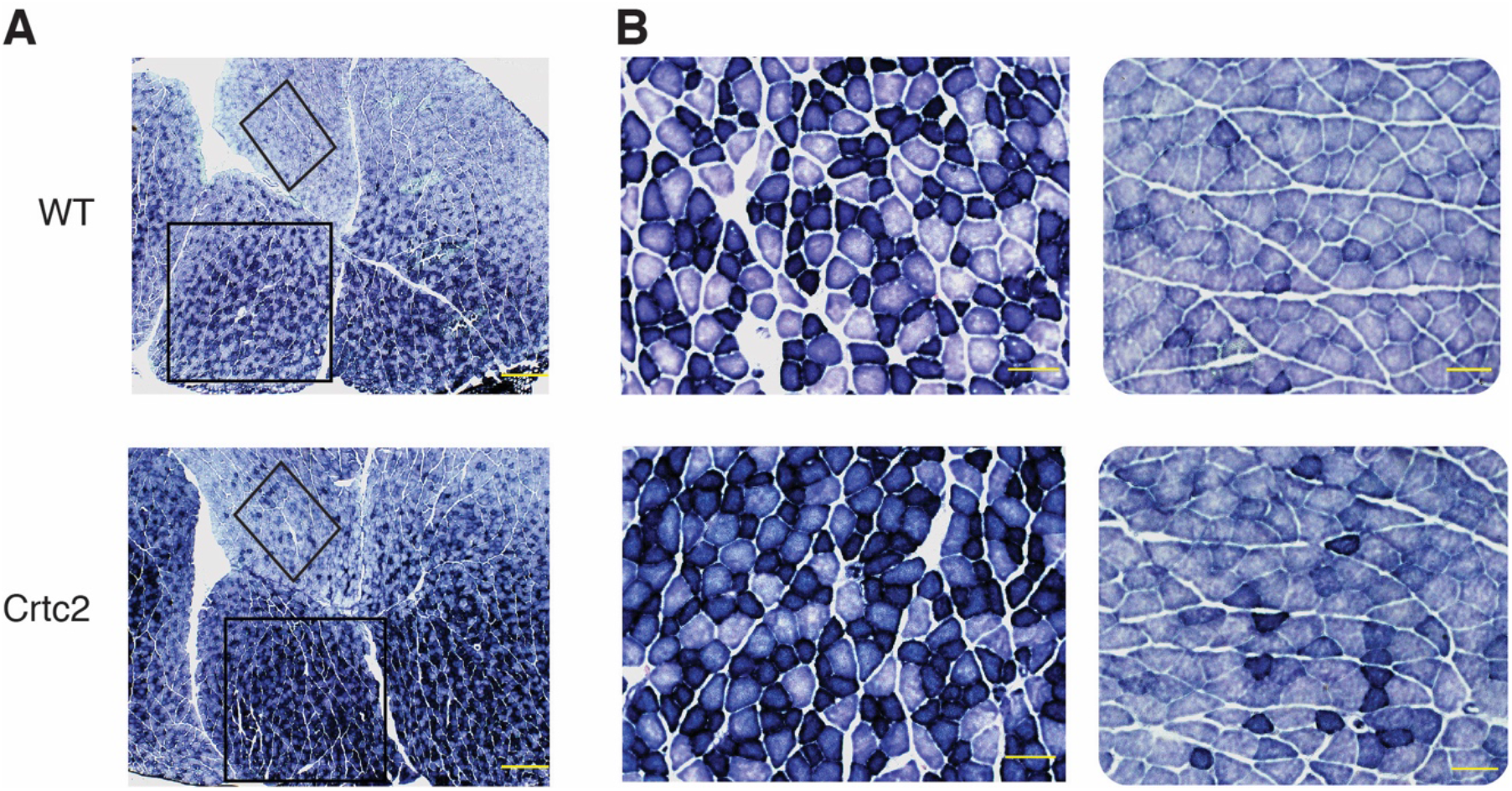
Analysis of succinate dehydrogenase in skeletal muscle fibers. (**A–CB**) Histological analysis of succinate dehydrogenase in myofibers from gastrocnemius muscle sections of WT and Crtc2 mice after Dox treatment. Boxes highlight areas of lighter and darker staining **B**) Scale bar = 2 mm. **C**) Scale bar = 100 μm. Close up from boxed areas in **B**)

**Supplemental Fig. 2.**
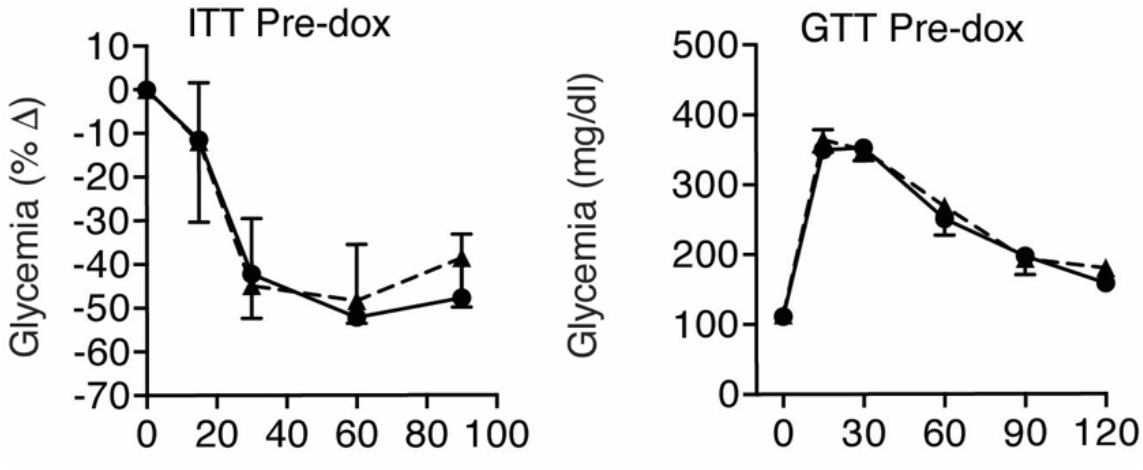
Glucose tolerance test controls. (**C–D**) GTT on WT and Crtc2 transgenic animals treated with dox for 1 week. Mice were fasted for 16 hrs. before i.p. injection of 20% glucose. n =10 mice per group.

**Supplemental Fig. 3.**
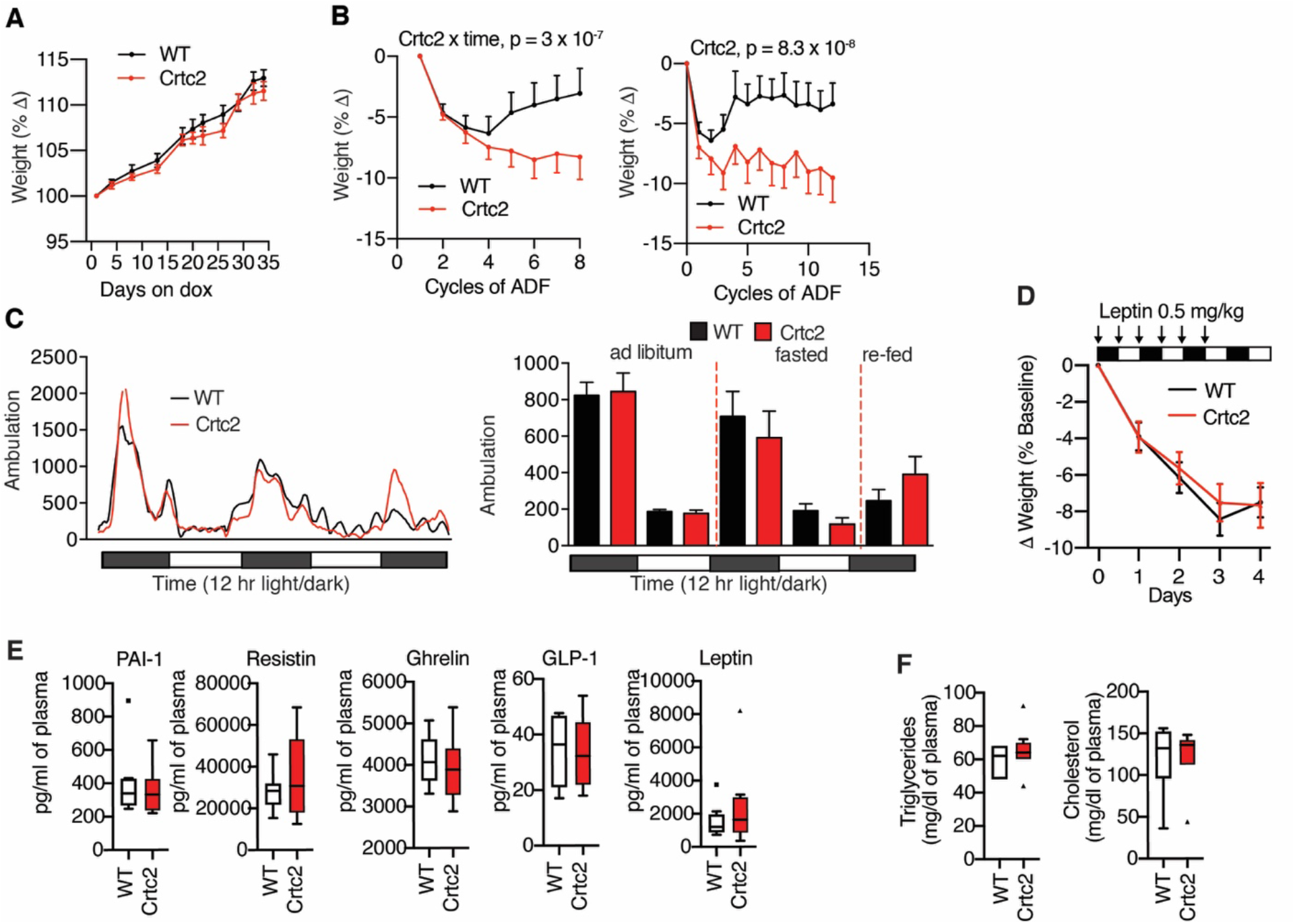
Effects of skeletal muscle overexpression of Crtc2 on the response to ADF and plasma feeding hormones in Crtc2 mice fed ad libitum. (**A**) Weight gain after treatment of 18-week old WT and Crtc2 transgenic mice (n=8) with doxycycline. (**B**) Changes in body weight during ADF. (**C**) Average ambulatory activity during ad libitum feeding, fasting and re-feeding, n = 8 mice per group. 12-hr averages of the data. There were no significant effects of Crtc2. **A–C)** Data shown as mean ± SEM. (**D**) WT or Crtc2 mice were injected twice daily with 0.5 mg/kg leptin and weight was monitored. *N*=8 mice per group. (**E–F**) Analysis of plasma proteins regulating feeding and triglycerides and cholesterol in Crtc2 expressing and WT mice.

**Supplemental Fig. 4.**
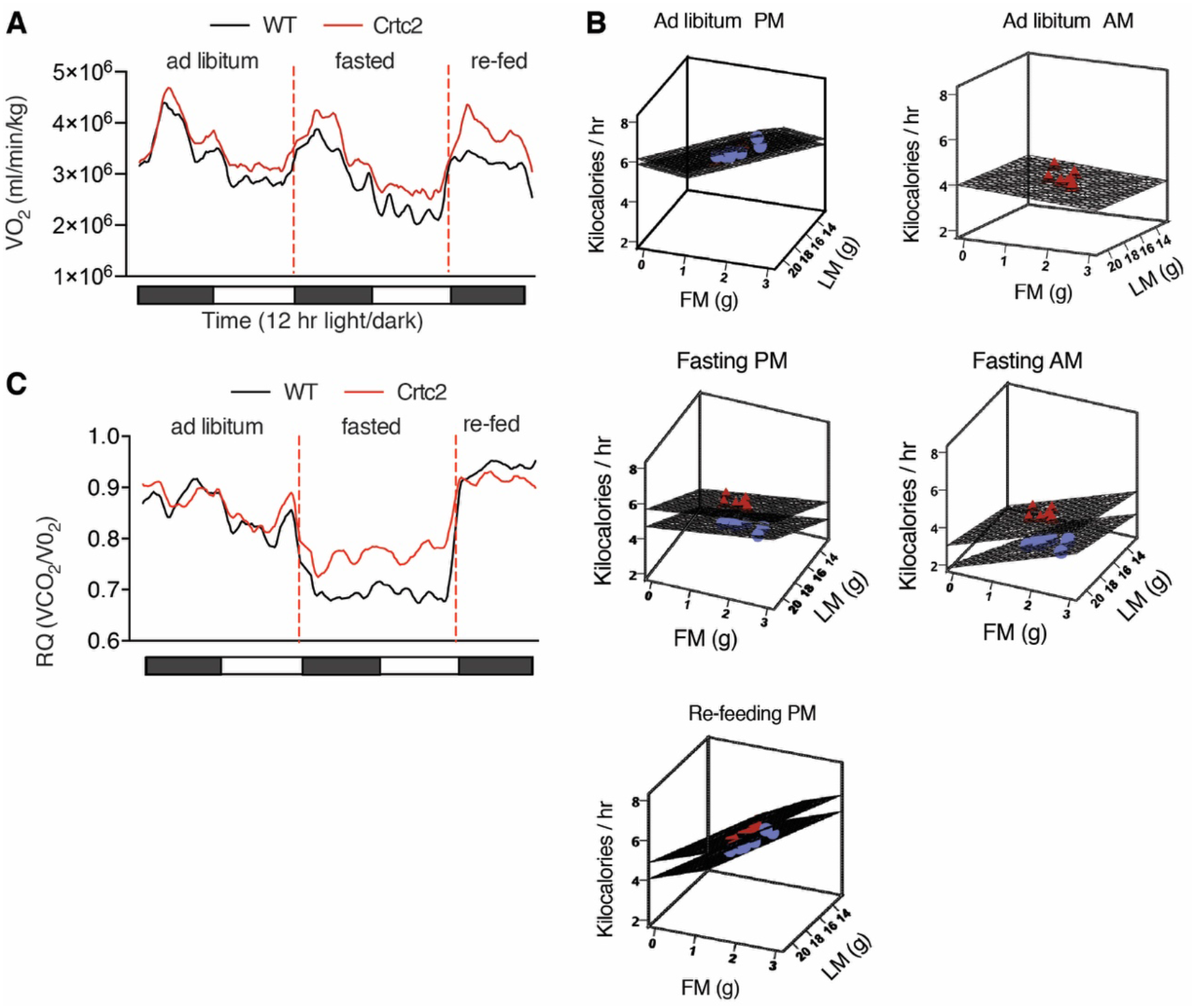
Whole-body energetics. (**A**) VO_2_ (and VCO_2_) were measured continuously for 72 hrs. in a CLAMS animal monitoring system during alternate day fasting. N = 8 per group (**B**) Total energy expenditure (EE) for WT and Crtc2 mice was calculated from VO_2_ and VCO_2_. The data was adjusted by repeated measures ANCOVA in with lean mass (LM) and fat mass (FM) as covariants at the following values: LM = 17.01 g; and FM = 1.62 g. Dot plot graphs shows adjusted means derived from repeated measure general linear model ANCOVA, using 5 consecutive 12-hr time intervals: PM-ad libitum; AM-ad libitum; PM-Fast; AM-Fast; and Re-feed EE as dependent variables, and LM and FM as covariates. The lack of intersection of the planes graphically demonstrates lack of covariation. (**C**) Respiratory Quotient from CLAMS experiment during alternate day fasting. Data are mean +SEM.

**Supplemental Fig. 5.**
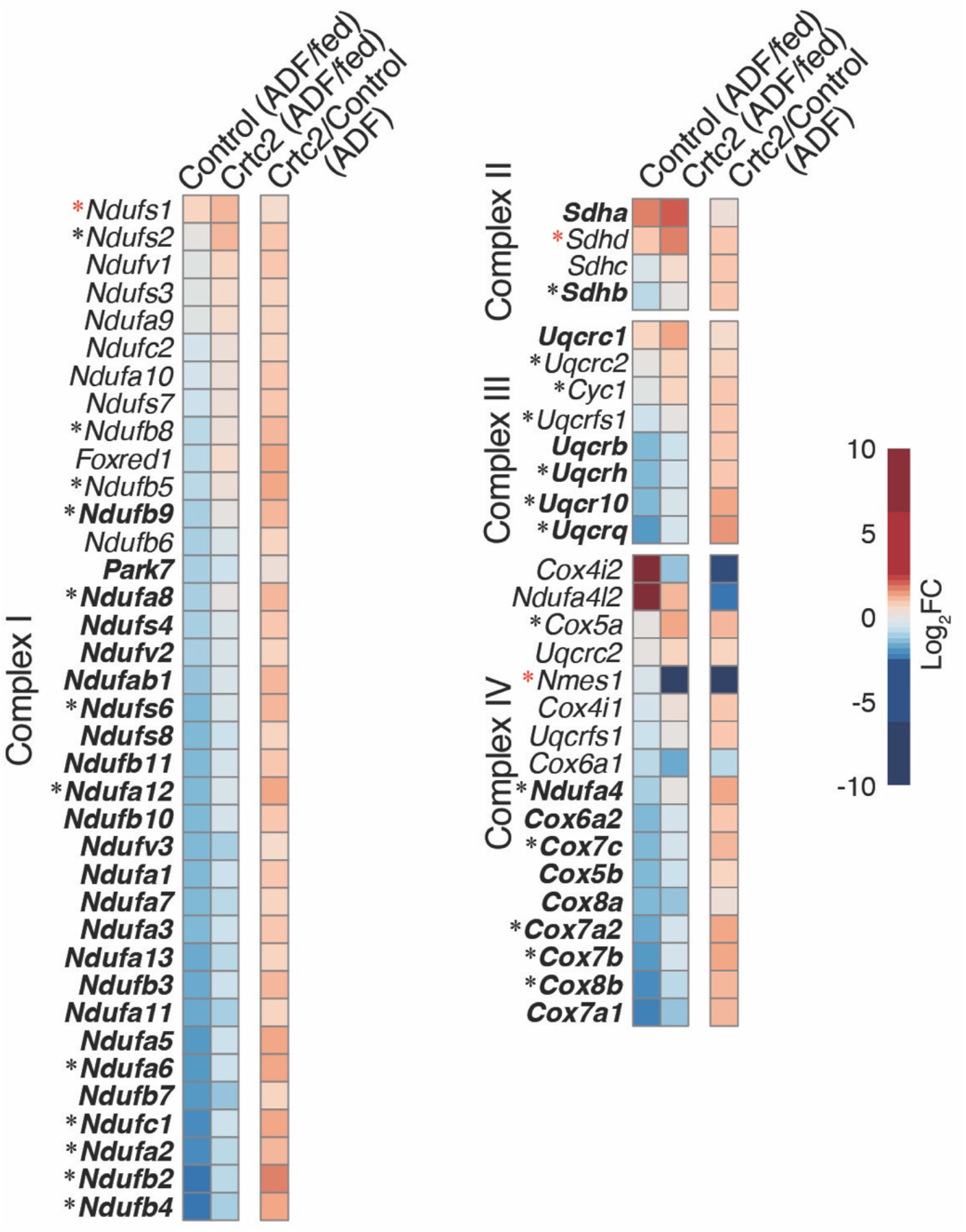
Electron transport chain gene expression. Expression profiles of genes that encode the electron transport chain. The mRNA levels in control and Crtc2-transduced TA muscles of mice subjected to ADF relative to the ad libitum fed mice (columns 1–2), and effect of Crtc2 transduction relative to control during ADF (column 3) are shown. ADF-regulated genes in control muscle appear in **bold**. *Crtc2-regulated genes in mice subjected to ADF. 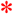ADF-regulated genes in Crtc2-transduced muscle but not control. FC, fold change

