## supplemental information for "Activation of the Crtc2/Creb1 transcriptional network in skeletal muscle enhances weight loss during intermittent fasting"

| Pathway name | # Entities found | # Entities total | FDR |
|---|---|---|---|
| Extracellular matrix organization | 45 | 329 | 3.98E-14 |
| Assembly of collagen fibrils and other multimeric structures | 17 | 67 | 2.12E-14 |
| MET promotes cell motility | 10 | 45 | 5.98E-07 |
| Elastic fibre formation | 12 | 46 | 6.13E-11 |
| Regulation of Insulin-like Growth Factor (IGF) transport and uptake | 13 | 127 | 2.47E-05 |
| Neutrophil degranulation | 26 | 480 | 2.35E-05 |
| Immune System | 79 | 2823 | 7.75E-06 |
| Signaling by Receptor Tyrosine Kinases | 23 | 622 | 7.13E-04 |
| Signaling by Interleukins | 25 | 639 | 0.049 |

**Supplemental Table 1. Gene pathway enrichment for secreted proteins among DEGs regulated by Crtc2 during ADF.**

**Supplemental Figures**

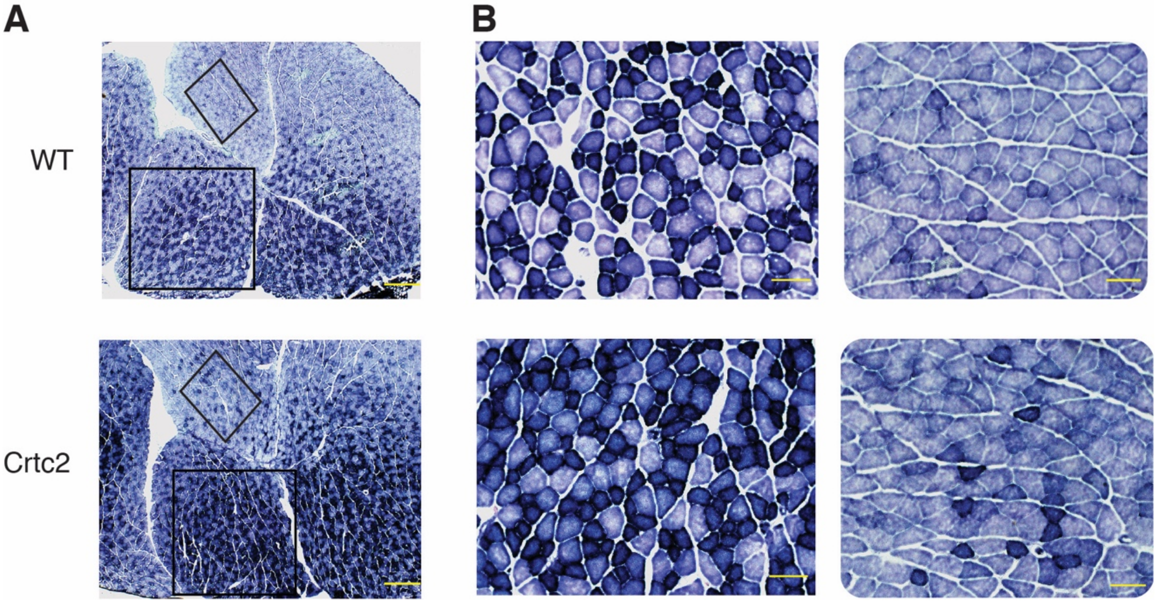


**Supplemental Fig 1. Analysis of succinate dehydrogenase in skeletal muscle fibers.**
(**A–CB**) Histological analysis of succinate dehydrogenase in myofibers from gastrocnemius muscle sections of WT and Crtc2 mice after Dox treatment. Boxes highlight areas of lighter and darker staining **B**) Scale bar = 2 mm. **C**) Scale bar = 100 μm. Close up from boxed areas in **B**)

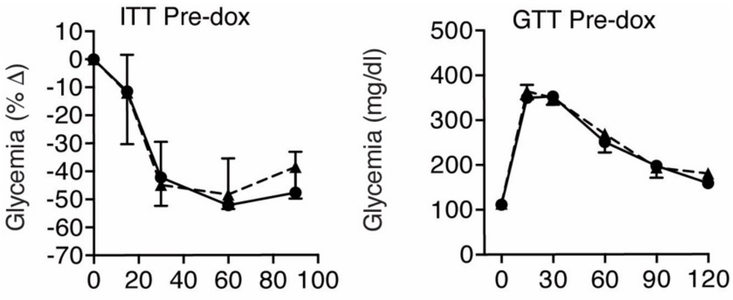


**Supplemental Fig. 2. Glucose tolerance test controls**

(**C–D**) GTT on WT and Crtc2 transgenic animals treated with dox for 1 week. Mice were fasted for 16 hrs. before i.p. injection of 20% glucose. n =10 mice per group.

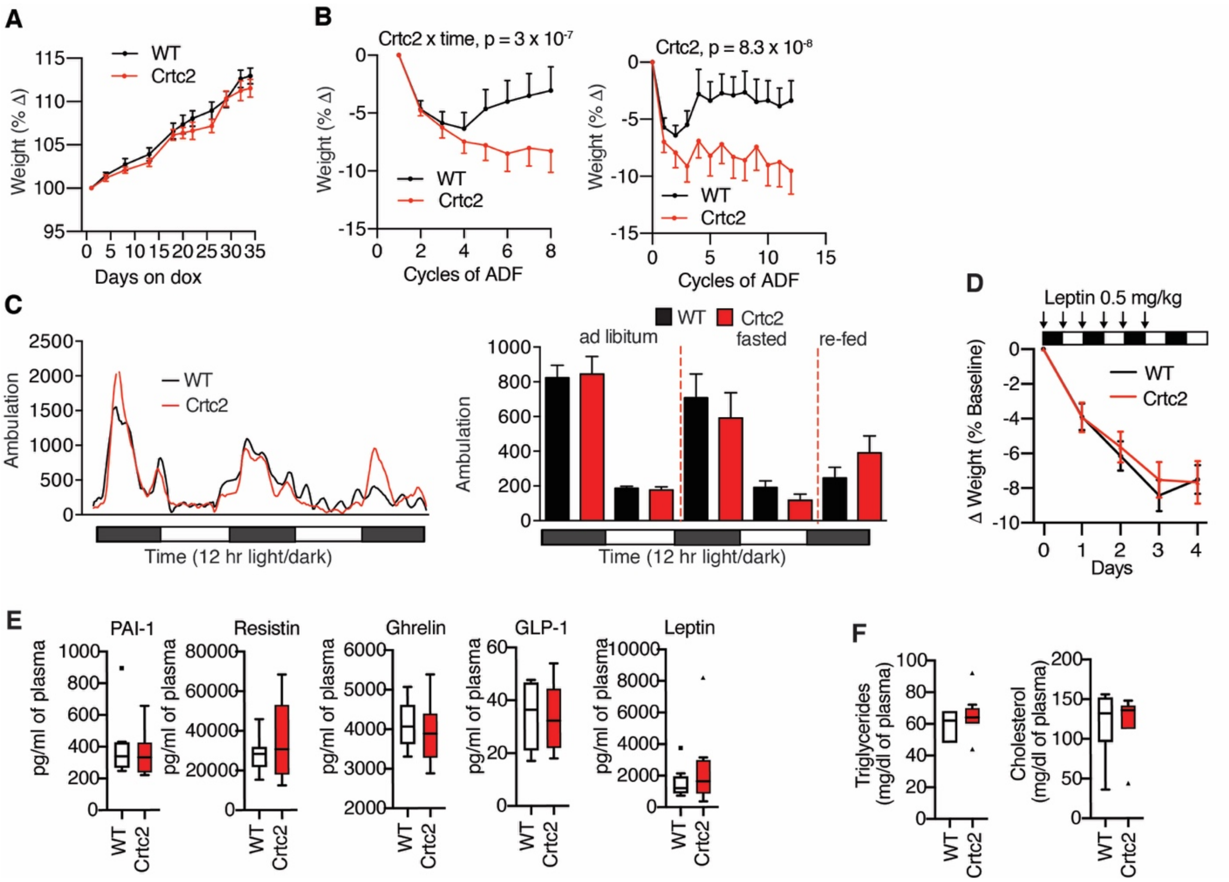


**Supplemental Fig. 3. Effects of skeletal muscle overexpression of Crtc2 on the response to ADF and plasma feeding hormones in Crtc2 mice fed ad libitum.**

(**A**) Weight gain after treatment of 18-week old WT and Crtc2 transgenic mice (n=8) with doxycycline.

(**B**) Changes in body weight during ADF.

(**C**) Average ambulatory activity during ad libitum feeding, fasting and re-feeding, n = 8 mice per group. 12-hr averages of the data. There were no significant effects of Crtc2. **A–C)** Data shown as mean ± SEM.

(**D**) WT or Crtc2 mice were injected twice daily with 0.5 mg/kg leptin and weight was monitored. *N*=8 mice per group.

(**E–F**) Analysis of plasma proteins regulating feeding and triglycerides and cholesterol in Crtc2 expressing and WT mice.

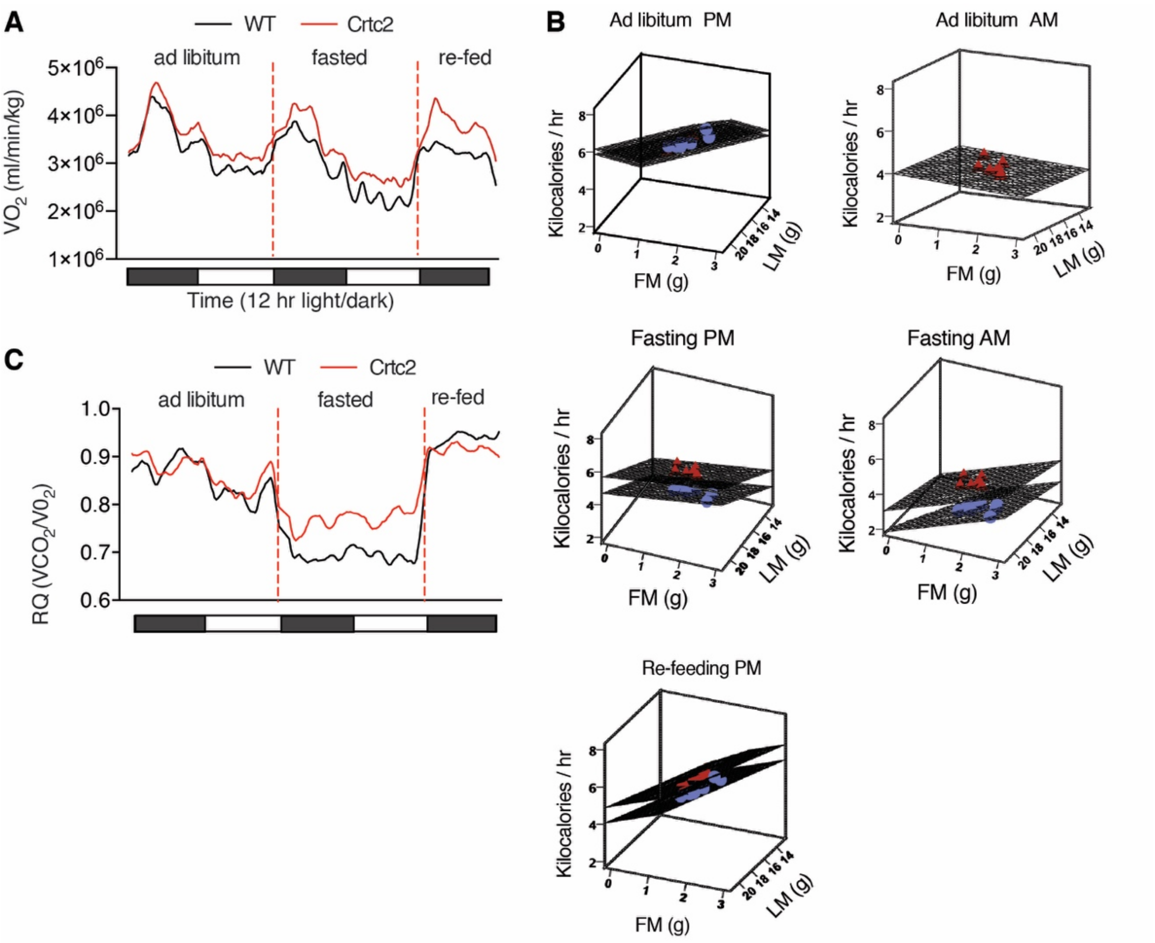


**Supplemental Fig. 4 Whole-body energetics.**

(**A**) $VO_2$ (and $VCO_2$) were measured continuously for 72 hrs. in a CLAMS animal monitoring system during alternate day fasting. N = 8 per group

(**B**) Total energy expenditure (EE) for WT and Crtc2 mice was calculated from $VO_2$ and $VCO_2$. The data was adjusted by repeated measures ANCOVA in with lean mass (LM) and fat mass (FM) as covariants at the following values: LM = 17.01 g; and FM = 1.62 g. Dot plot graphs shows adjusted means derived from repeated measure general linear model ANCOVA, using 5 consecutive 12-hr time intervals: PM-ad libitum; AM-ad libitum; PM-Fast; AM-Fast; and Re-feed EE as dependent variables, and LM and FM as covariates. The lack of intersection of the planes graphically demonstrates lack of covariation.

(**C**) Respiratory Quotient from CLAMS experiment during alternate day fasting. Data are mean +SEM.

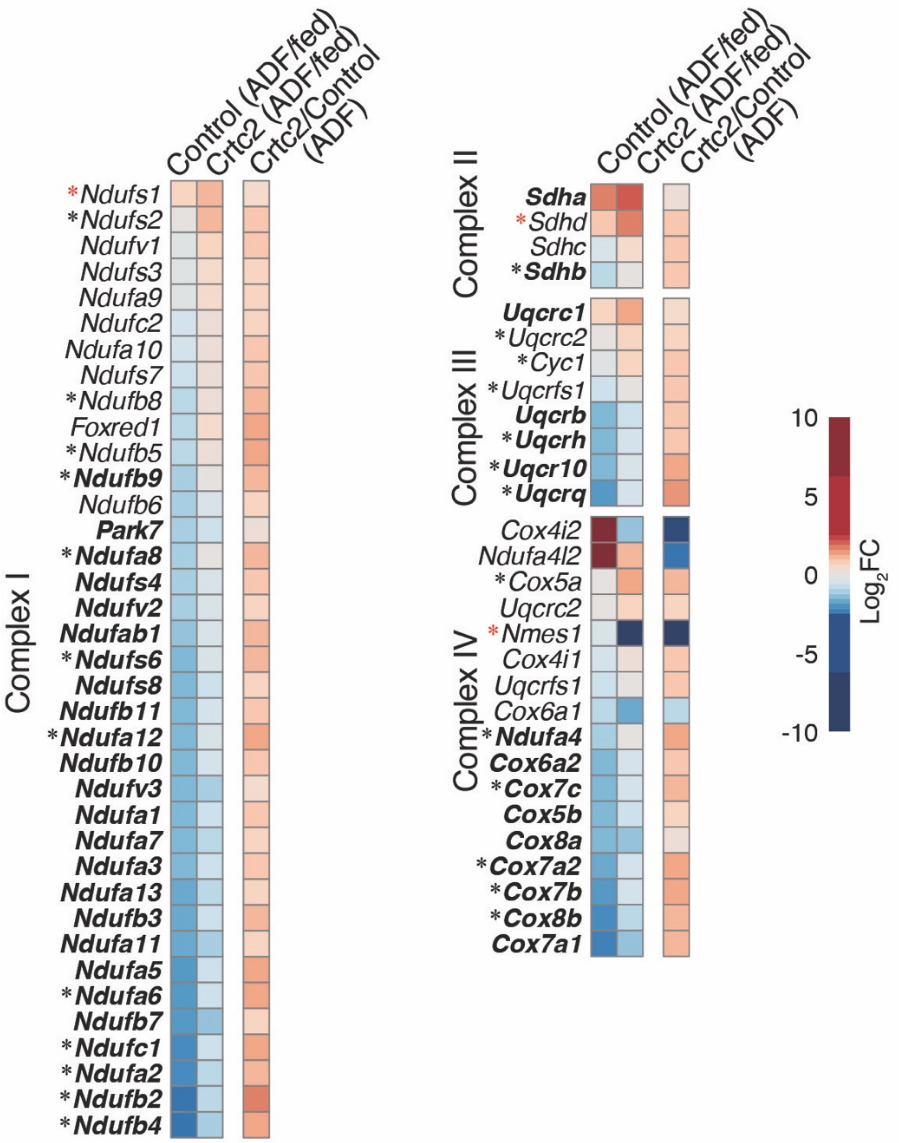


**Supplemental Fig. 5 Electron transport chain gene expression**

Expression profiles of genes that encode the electron transport chain. The mRNA levels in control and Crtc2-transduced TA muscles of mice subjected to ADF relative to the ad libitum fed mice (columns 1–2), and effect of Crtc2 transduction relative to control during ADF (column 3) are shown. ADF-regulated genes in control muscle appear in **bold**. *Crtc2-regulated genes in mice subjected to ADF. *ADF-regulated genes in Crtc2-transduced muscle but not control. FC, fold change
